# Spindle scaling is governed by cell boundary regulation of microtubule nucleation

**DOI:** 10.1101/2020.06.15.136937

**Authors:** Elisa Maria Rieckhoff, Frederic Berndt, Stefan Golfier, Franziska Decker, Maria Elsner, Keisuke Ishihara, Jan Brugués

**Affiliations:** Max Planck Institute of Molecular Cell Biology and Genetics, 01307 Dresden, Germany; Max Planck Institute for the Physics of Complex Systems, 01187 Dresden, Germany; Centre for Systems Biology Dresden, 01307 Dresden, Germany; Cluster of Excellence Physics of Life, TU Dresden, 01062 Dresden, Germany

## Abstract

Cellular organelles such as the mitotic spindle adjust their size to the dimensions of the cell. It is widely understood that spindle scaling is governed by regulation of microtubule polymerization. Here we use quantitative microscopy in living zebrafish embryos and *Xenopus* egg extracts in combination with theory to show that microtubule polymerization dynamics are insufficient to scale spindles and only contribute below a critical cell size. In contrast, microtubule nucleation governs spindle scaling for all cell sizes. We show that this hierarchical regulation arises from the partitioning of a nucleation inhibitor to the cell membrane. Our results reveal that cells differentially regulate microtubule number and length using distinct geometric cues to maintain a functional spindle architecture over a large range of cell sizes.

## Main Text

During early animal development, a succession of rapid cell divisions decreases cellular volumes in the absence of embryonic growth. These reductive cell divisions require that intracellular structures scale with cell size to ensure that proportionality is maintained (*1*-*3*). One essential structure that needs to scale to maintain the robust segregation of chromosomes during early embryogenesis is the mitotic spindle. Spindles can scale an order of magnitude and fall into what appears to be a universal scaling relationship with cell size (*4*-*20*). Despite this phenomenological characterization, however, we still lack a mechanistic understanding of how microtubule-based processes regulate spindle size relative to cell size.

Current models propose that spindles scale by changing the length of microtubules via a limiting cellular component that regulates microtubule polymerization dynamics (*10, 11, 17, 21, 22*). Consistent with this proposal, microtubule growth velocity correlates with spindle size in early *C. elegans* and sea urchin embryos (*17*), and increased activity of the microtubule polymerase XMAP215 is sufficient to change spindle length *in vitro* and *in vivo* (*16, 22*). These studies on spindle size have been limited to measurements of microtubule dynamics in relatively small spindles (spindle length L < 20 µm) or in biochemically perturbed large spindles *in vitro* (L ∼ 50 µm). However, detailed quantitative measurements of microtubule dynamics in a living embryo that covers the whole range of spindle scaling are missing (*12*). Consequently, how microtubule dynamics are regulated during the dramatic changes in spindle length during animal development (L = 5 - 50 µm) is unknown.

Here, we systematically quantify microtubule dynamics, nucleation and organization in spindles over the entire range of cell sizes in early zebrafish embryos. We show that, in contrast to previous models, spindle size is scaled by microtubule nucleation across the entire range of cell sizes. In contrast, microtubule dynamics only influences scaling in small cells, but remains insufficient to account for spindle scaling. Our data is consistent with a theory in which component limitation of microtubule nucleators and membrane partitioning of a nucleation inhibitor quantitatively explain both the exact scaling of spindles with cell size and also the hierarchical regulation of microtubule nucleation and dynamics. In agreement with this theory, when spindles are assembled without cell membranes in encapsulated *Xenopus* egg extracts, spindles scale exclusively by microtubule nucleation. We propose that this hierarchical regulation is key to maintaining proper microtubule density and organization in spindles as they dramatically change in size during development.

## Results

### Changes in microtubule dynamics are insufficient to account for spindle scaling

To simultaneously characterize spindle scaling, microtubule dynamics, and cell volume, we used early zebrafish embryos as a model system. Their large size (∼700 µm in diameter), rapid development, and optical transparency, make zebrafish embryos an ideal system to study spindle scaling over three orders of magnitude in cell size (*23*). Using light-sheet microscopy, we imaged spindles and cell boundaries in the entire zebrafish embryo during the first 5 hours of development, Fig 1A and Movie S1. While previous studies were mainly restricted to measuring spindle length (*4, 8*-*12, 17, 22, 24*), our approach allowed us to segment spindle and cellular volumes in 3D and obtain the volumetric relationship between spindle size and cell size (FigS1 and Movie S2). We found that during early zebrafish embryogenesis, spindle and cell volumes decrease 10- and 1000-fold, respectively, covering most of the range of spindle scaling measured in other species (*12*) (Fig 1B-C and Fig S1). We found that spindle scaling in zebrafish embryos exhibits two distinct regimes, similar to *Xenopus laevis* embryos (*4*). In large cells, spindle size saturates above a critical cell size (cell volume V_cell_ > 10^6^ µm^3^, cell diameter d_cell_ > 125 µm) and becomes independent of cell size (‘upper limit’), whereas below this critical cell size, it scales with cell size (‘scaling regime’, see Fig 1B-C).

**Fig. 1:**
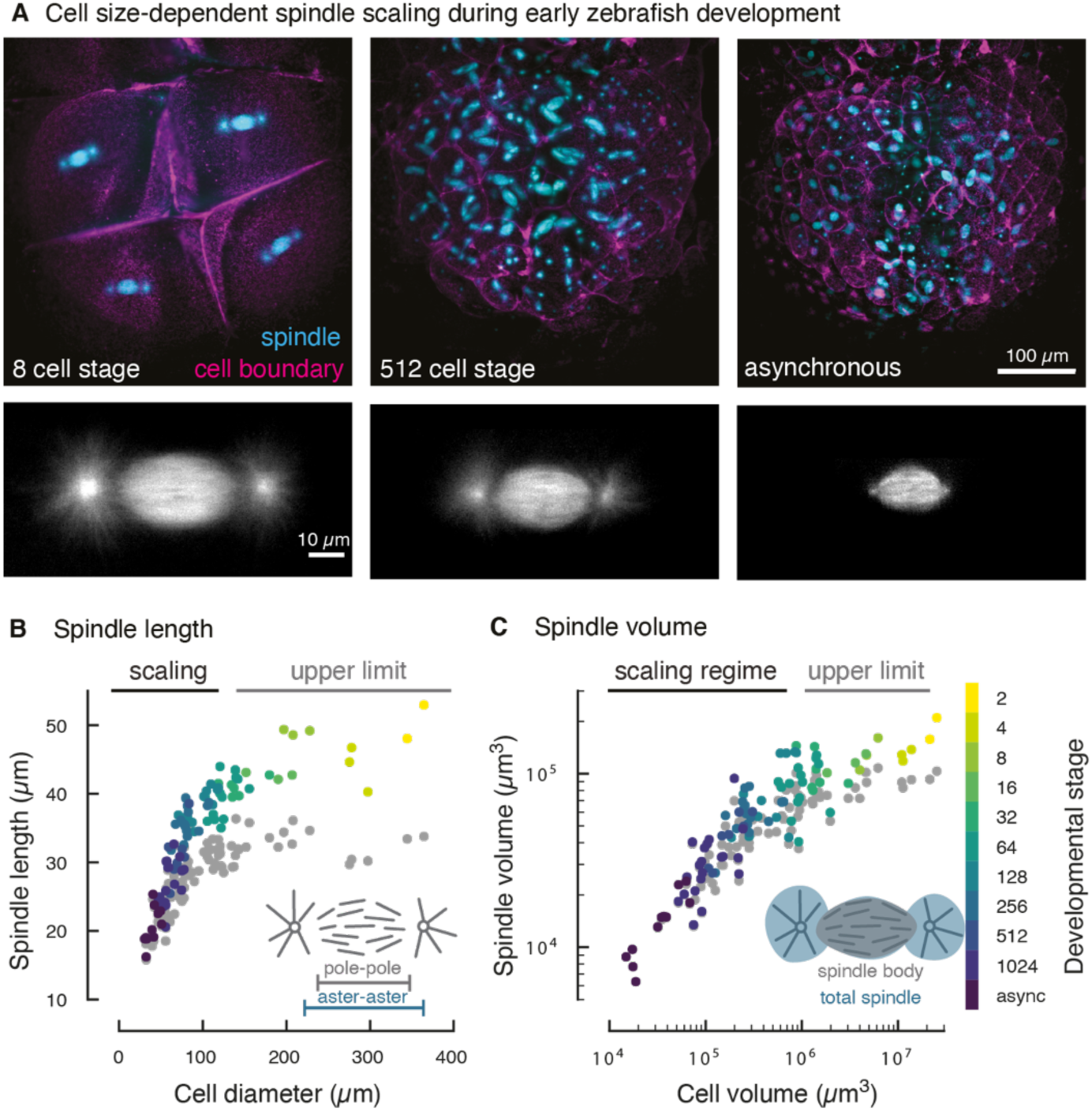
Cell size-dependent spindle scaling during early zebrafish development. (A) 3D light-sheet microscopy of spindles and cell boundaries in zebrafish embryos. Maximum intensity projections of a fluorescently labeled zebrafish embryo at selected developmental stages (see Movie S1). (B) Spindle scaling in early zebrafish embryos covers two regimes. In large cells, spindle length remains constant (‘upper limit’). Below a critical cell diameter (d_cell_ = 125 µm), spindle length scales linearly with cell diameter. Each dot denotes an individual measurement of aster-to-aster spindle length (color-coded by developmental stage) or pole-to-pole spindle length (in gray, 96 analyzed spindles and cells in 4 different embryos). (C) 3D segmentation of spindles and cells (see Fig S1) yields the scaling relationship between spindle volume and cell volume. The scaling of the total spindle volume (including asters, color-coded by cell stage) and the spindle body volume (excluding asters, gray) is nearly indistinguishable.

To characterize the contribution of microtubule dynamics to spindle scaling, we analyzed the polymerization velocity (v_p_), the depolymerization velocity (v_d_), and microtubule lifetime (*τ*). To measure microtubule polymerization velocity, we microinjected fluorescently labeled microtubule end-binding proteins (EB1-GFP) (*25, 26*) into zebrafish embryos at the one-cell stage (Fig 2A and Movie S3). We acquired time-lapse movies during the subsequent embryonic cell divisions and tracked growing microtubule plus ends (see Materials and Methods and Fig S4A-E). We found that microtubule polymerization velocities scale linearly with spindle length for spindles shorter than ∼30 µm. Strikingly, polymerization velocities plateau for larger spindles and become independent of spindle size with an average velocity v_p_ = 25 ± 2 µm/min (Fig 2A). In this regime of constant microtubule polymerization velocity, spindles still increase in length approximately two-fold from ∼30 µm to 50 µm. The correlation we observed between microtubule polymerization velocity and cell size in small zebrafish cells is consistent with recent studies in *C. elegans* and sea urchin spindles (*17*). However, the 1.4-fold change in microtubule growth velocity in this regime is significantly lower than the corresponding 5-fold change in spindle length (Fig 2A). Moreover, microtubule polymerization velocity plateaus before spindle size saturates. These observations together indicate that the changes in microtubule polymerization velocity alone cannot account for the observed scaling of mitotic spindles.

**Fig. 2:**
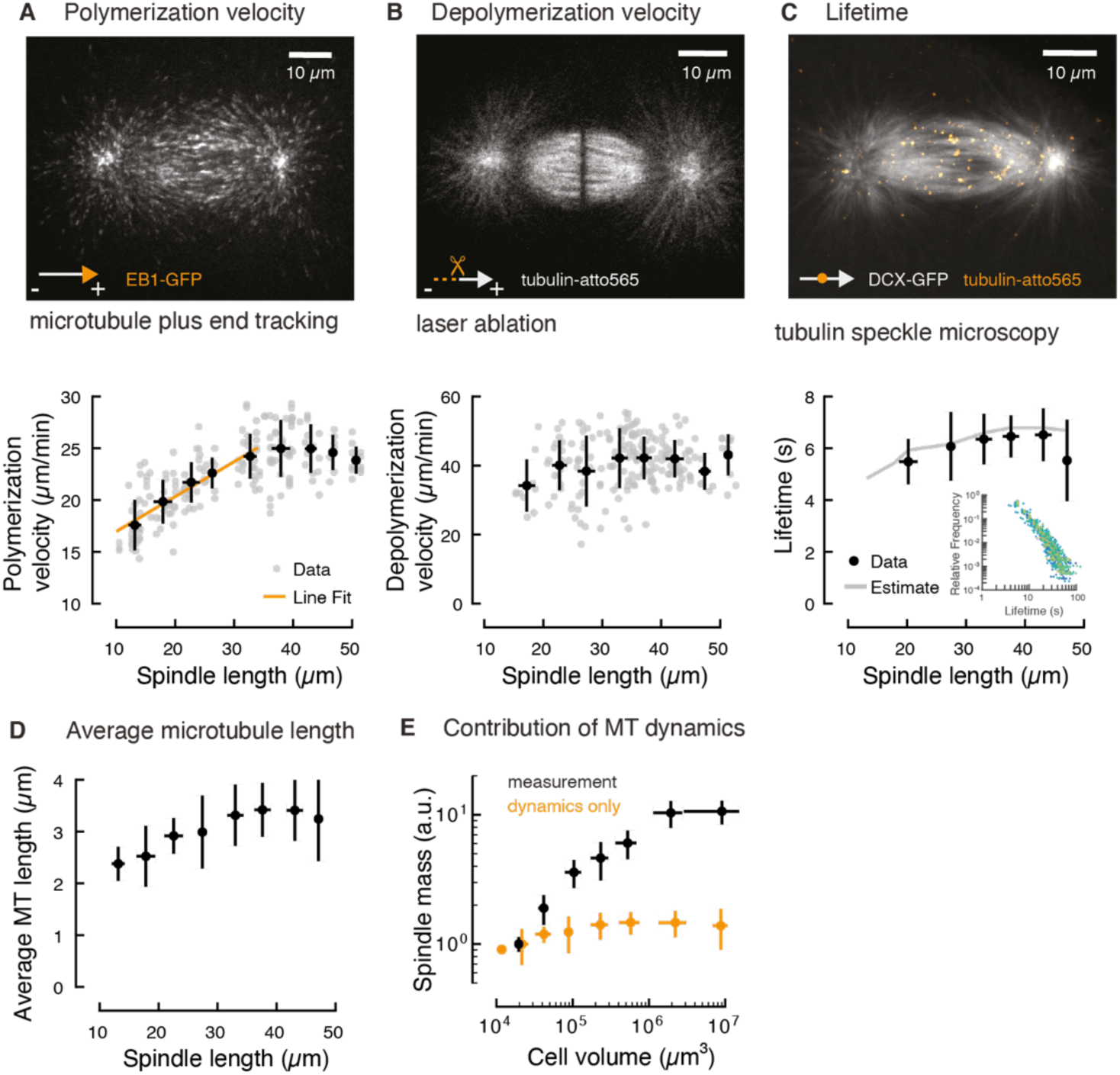
Microtubule dynamics are insufficient to account for spindle scaling. (A) Quantification of microtubule polymerization in spindles of different sizes during early zebrafish development shows two regimes, see also Fig S4 and Movie S3. Microtubule growth velocity scales linearly with spindle length in spindles smaller than 35 µm. In larger spindles, microtubule polymerization velocity remains constant. Individual and binned measurements are shown in gray (N=196) and black respectively (mean ± SD, bin size 5 µm). (B) Laser ablation reveals that microtubule depolymerization velocity is independent of spindle length, see also Fig S2 and Movie S4 (N=202). (C) Tubulin speckle microscopy reveals that microtubule lifetimes are constant in spindles of different sizes, see also Fig S5 and Movie S5. Solid gray line is the estimate derived from changes in the microtubule polymerization velocity observed during scaling (Materials and Methods). Inset shows the measured speckle lifetime distributions color-coded by spindle length (N = 47). (D) Microtubule length correlates with spindle size in spindles smaller than 35 µm, but remains constant in larger spindles. Average microtubule length obtained from the measured microtubule dynamics (Materials and Methods). Error bars are obtained from error propagation of the measurements of microtubule dynamics. (E) Modulation of microtubule dynamics is not sufficient to account for spindle scaling. The estimated scaling of spindle mass from microtubule dynamics alone is significantly lower than the measured spindle mass (in black).

We next measured microtubule depolymerization velocities in differently sized spindles by severing fluorescently labeled microtubules using femtosecond laser ablation (*15, 27, 28*). Laser ablation induces synchronous waves of microtubule depolymerization that can be tracked over time to measure microtubule depolymerization velocity (see Materials and Methods, Fig S2 and Movie S4). Using this method, we found that microtubules depolymerize at 41 ± 7 µm/min, independently of spindle size (Fig 2B). Thus, microtubule depolymerization velocity is not regulated by cell size and does not contribute to spindle scaling.

Finally, to investigate potential variations in microtubule turnover during scaling, we quantified microtubule lifetimes across different spindle sizes using tubulin speckle microscopy (*29*-*31*). We found that the measured lifetime distributions in spindles of lengths ranging from 15 µm to 50 µm fall onto a single curve with average turnover rate of 6 ± 2 s, Fig 2C, Fig S5 and Movie S5. Thus, microtubule lifetimes do not change significantly over a large range of spindle sizes.

We used these three measurements of microtubule dynamics to estimate the average length of microtubules during spindle scaling (see Materials and Methods). These results show that microtubules in zebrafish spindles are significantly shorter than the overall spindle length. For large spindles, constant microtubule dynamics imply that microtubule length plateaus at an average of 3.3 ± 0.8 µm. In smaller spindles, microtubule length correlates with spindle length (Fig 2D). This scaling of microtubule length is also consistent with laser ablation measurements (Fig S2G). However, microtubule length varies only slightly from 3.3 µm to 2.3 µm in spindles ranging from 35 µm to 10 µm in length, respectively. The scale-invariant microtubule length in large spindles together with the subtle change in microtubule dynamics in small cells suggests that microtubule dynamics alone are insufficient to account for spindle scaling.

To explore the contribution of microtubule dynamics to the scaling of spindle mass, we estimated the change in spindle mass for different cell sizes based solely on the measured change in microtubule length. The total microtubule mass in a spindle M_spd_ is proportional to the microtubule number N_MT_ times the average microtubule length L_MT_, *M*_*spd*_ = *m N*_*MT*_,*L*_*MT*_, where m is the average microtubule mass per unit length. Under the assumption that microtubule number is constant—to account only for the contribution of microtubule dynamics—, we find that the contribution of microtubule dynamics to the change in spindle mass is drastically lower compared to the measured spindle mass (Fig 2E). Consequently, we conclude that changes in microtubule dynamics are insufficient to account for spindle scaling.

### Changes in microtubule number determine spindle scaling

The surprisingly negligible effect of microtubule dynamics on spindle scaling is at odds with previous models. Instead, it suggests that microtubule number must scale with cell size to regulate spindle size. To test for this possibility, we quantified the change in microtubule number during spindle scaling using two independent approaches. First, we used our measurements of spindle mass and microtubule length to infer how microtubule number must change during spindle scaling (Fig 3A). Second, we directly measured the microtubule number via laser ablation experiments (see Materials and Methods and Supplemental Information). Using these two independent methods, we found that microtubule number decreases approximately 10-fold during spindle scaling in early zebrafish embryos (Fig 3A). Thus, small spindles contain significantly fewer microtubules than large spindles. Altogether, these data show a hierarchical regulation of processes that control spindle scaling (Fig 3A). Large spindles adjust their size exclusively by tuning microtubule number, while microtubule length remains unchanged. As spindles become smaller, regulation of microtubule length begins to have a limited contribution but remains insufficient to account for spindle scaling.

**Fig. 3:**
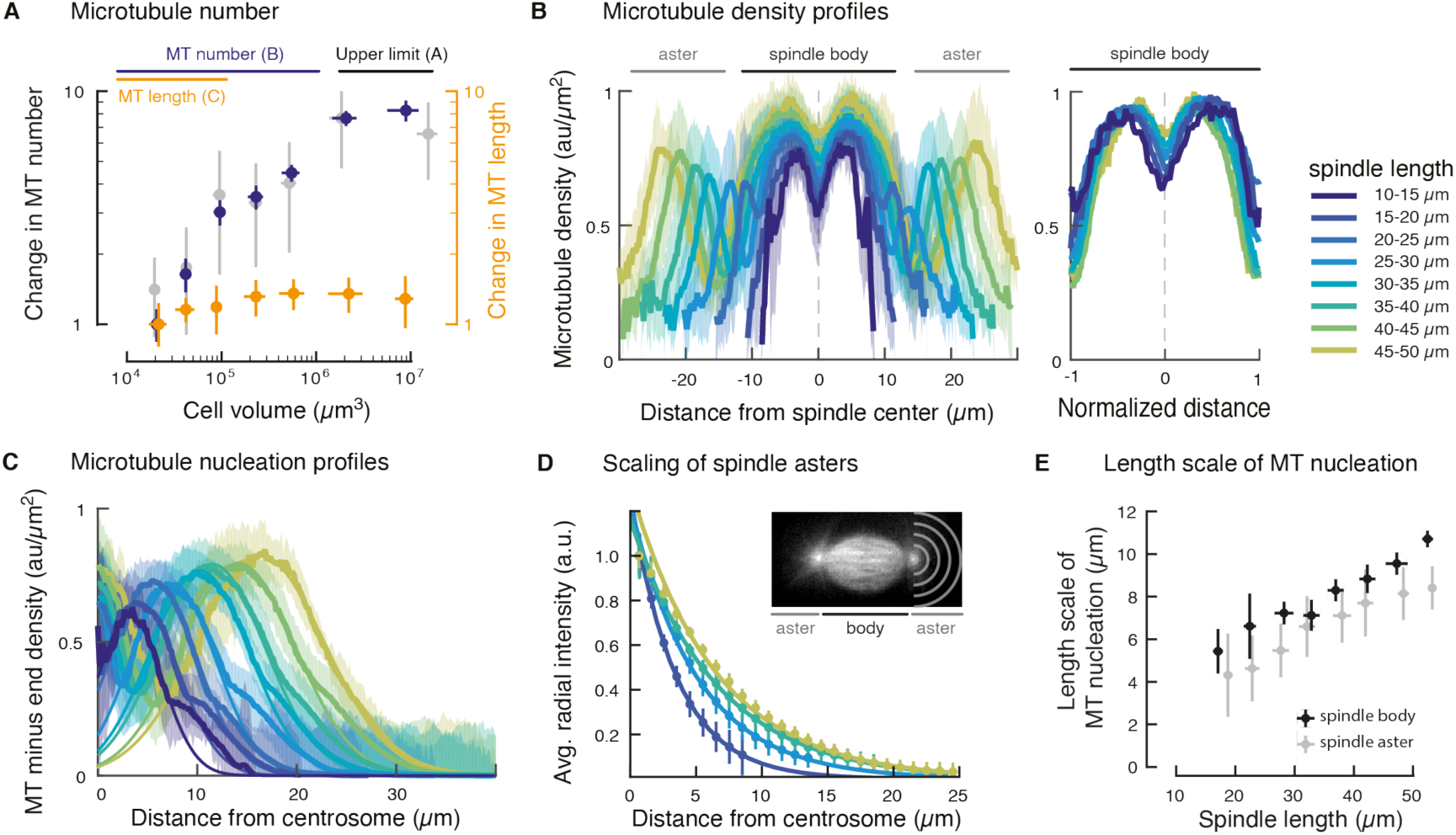
Microtubule nucleation determines spindle scaling and architecture. (A) The number of microtubules within the spindle decreases 10-fold during spindle scaling. Microtubule number N_MT_ was calculated from measurements of spindle mass M_spd_ and microtubule length L_MT_ (N_MT_ = M_spd_ / L_MT_, in blue), and laser ablation (in gray, see Supplemental Information). For comparison, the change in microtubule length during spindle scaling is shown in orange. Error bars denote standard deviations obtained from error propagation. (B) Left, microtubule density profiles scale with spindle size. Right, rescaled microtubule density profiles in the spindle body are shape-invariant (N = 210). Shaded regions denote standard deviations. (C) Microtubule nucleation profiles for microtubules of the same polarity show that nucleation occurs throughout the spindle. Thin solid lines correspond to Gaussian fits to estimate the length scale of microtubule nucleation. (D) Spindle asters scale with spindle length. The size of spindle asters was determined by radially averaging the tubulin fluorescence intensity at different distances from the centrosome (denoted by gray half circles, N = 165). The normalized radial fluorescence intensities decay exponentially with distance from the centrosome–indicative of non-centrosomal microtubule nucleation (solid lines). Error bars correspond to standard deviations. (E) The extent of microtubule nucleation in the spindle body (in black) and in the asters (in gray) decreases linearly with spindle length.

### Scaling of microtubule nucleation profiles governs spindle size and architecture

Our data suggests that modulating microtubule number in spindles is the primary mechanism for spindle scaling. Microtubule number in spindles is regulated by the extent of microtubule nucleation. To gain further insight into microtubule nucleation during scaling, we investigated its spatial distribution in spindles of different sizes using laser ablation. Briefly, analysis of laser ablation-induced microtubule depolymerization reveals the microtubule minus end density at the ablation site. In the absence of significant microtubule fluxes, the minus end density reveals the location and amount of microtubule nucleation (*15*). In zebrafish spindles, microtubule transport occurs at a velocity of 3.4 ± 1.6 µm/min (Fig S4H). This velocity together with the average microtubule lifetime of 6 s, implies that microtubules move less than 0.5 µm away from their nucleation site, which is negligible compared to the size of these spindles. Thus, in our system, the density profile of microtubule minus ends is a good approximation of the profile of microtubule nucleation.

By analyzing laser cuts performed at different positions within the spindle, we found that microtubule minus end density is roughly constant within the spindle body, with a dip at the spindle center due to volume exclusion by DNA (*12, 22*) (see Supplemental Information, Fig 3B). These minus end density profiles can only be maintained by non-centrosomal microtubule nucleation occurring throughout the spindle (*14, 15, 32*-*35*). Consistent with this, we found that the nucleation profiles of microtubules of the same polarity spread throughout the spindle (Fig 3D and Fig S3). Additionally, the profiles of centrosomal microtubules decay more slowly than the inverse of the distance (1/R) that would be predicted by purely centrosomal nucleation (*32*) (Fig 3C). The spatial extent of microtubule nucleation within the spindle and away from centrosomes scales linearly with spindle size (Fig 3E). These nucleation profiles can be collapsed into a single profile by rescaling space (Fig 3B), suggesting that their shape is invariant to cell size. Taken together, our data suggests that the scaling of the spatial microtubule nucleation profiles leads to a reduced number of microtubules in smaller spindles, while its invariant shape ensures that the same spindle architecture is maintained across all sizes (Fig S3).

### A model based on membrane sequestration of a microtubule nucleation inhibitor accounts for hierarchical spindle scaling

Previous studies proposed that cell volume regulates spindle size by providing a limiting pool of cytoplasmic components required for spindle assembly (*10, 11, 36*). Our quantitative measurements of microtubule number and length show that spindles mainly scale by modulating microtubule nucleation. Consequently, factors affecting microtubule nucleation (referred to as nucleators) must become limiting first. Because activation of nucleators happens in the proximity of chromosomes (*14, 15*), this activation process may be limited by diffusion when cells are large enough (*37*).

In the simplest limiting component model, nucleators would scale with cell volume and saturate beyond the diffusion-limited volume (Fig 4A and Supplemental information), which qualitatively agrees with our data (regime A to B). The exact scaling behavior of microtubule number N_MT_ can be analytically solved and is well captured by the expression 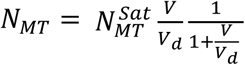, where V is cell volume and V_d_ is the diffusion-limited volume (see Supplemental information). However, linear scaling of microtubule number with cell volume poses a problem for the limited scaling of microtubule dynamics in small cells. In a limiting component model, factors regulating microtubule polymerization dynamics (e.g. tubulin, polymerases) would also scale linearly with cell volume, implying that the ratio of microtubule number and components that regulate microtubule dynamics would remain constant (Fig 4B). Thus, if microtubule nucleation is limiting first, then we should expect components regulating microtubule polymerization dynamics to never become limiting. As a consequence, either microtubule number or components that regulate microtubule dynamics need to scale differently with cell volume.

**Fig. 4:**
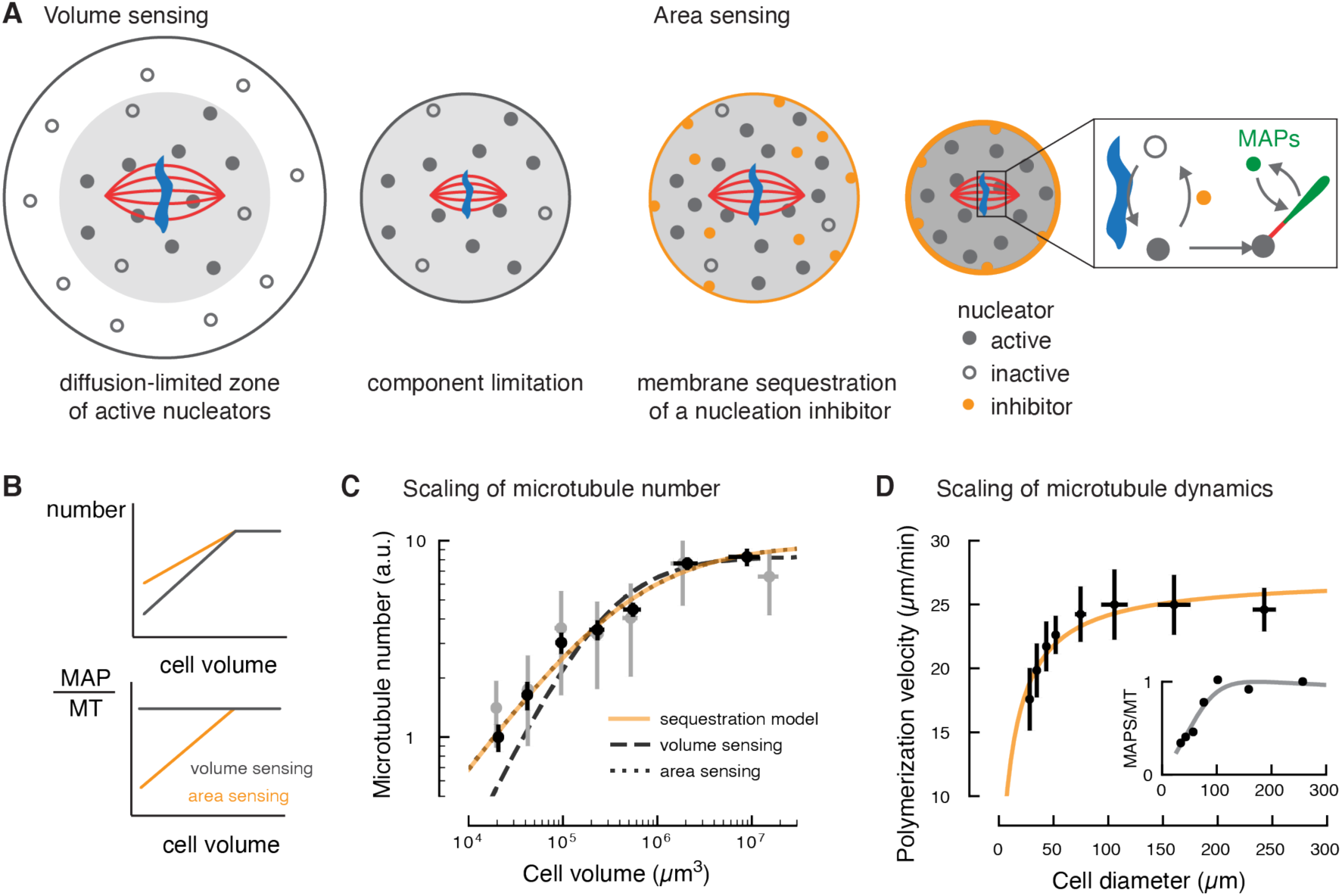
A theoretical model based on membrane sequestration of a microtubule nucleation inhibitor accounts for spindle scaling. (A) Left, schematic representation of the limiting component model (‘volume sensing’). In large cells, the activation of nucleators (shown as filled circles) near chromosomes is limited by diffusion. As cell size decreases below the diffusion-limited volume, the number of nucleators scale with cell volume. Right, schematic representation of the membrane sequestration model (‘area sensing’). An inhibitor of microtubule nucleation (orange circle) is adsorbed at the cell boundary. Inset: Activation of nucleators occurs around DNA (in blue), while an inhibitor of microtubule nucleation (orange) triggers its inactivation. Active nucleators can generate new microtubules (red line), to which microtubule-associated proteins (MAPs, in green) can bind. (B) Prediction of how volume sensing and area sensing affect microtubule number and dynamics. If both microtubule (MT) number and factors promoting microtubule growth (MAPs) scale equally with cell volume, their ratio remains constant. Only if they scale differently with cell size, their ratio changes and microtubule dynamics may change. (C) Microtubule number is sensitive to the surface area of the cell and it is accounted by the membrane sequestration model (see Supplemental Information, orange line). (D) The membrane sequestration model (orange line) captures the change in polymerization velocity with cell size (see Supplemental Information). Inset: The number of MAPs per microtubule (MT) decreases with decreasing cell diameter.

To explore the exact scaling of microtubule number with cell size, we fitted our measurements of microtubule number to the expression 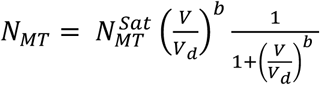, which captures the diffusion-limited regime but allows for a scaling that differs from the cell volume via the exponent b (b=1 corresponding to pure volume scaling). We found that the best fit for the scaling exponent is close to 2/3 (b = 0.64 ± 0.20), implying that microtubule number scales with the area of the cell (*V*^2/3^) and not with cell volume.

Our results suggest that microtubule number can sense the cell surface even though the activation of nucleation is a process that occurs mainly within the cytoplasm. One mechanism that allows microtubule nucleation to be coupled to the cell surface is the partitioning of a regulatory component into the cell membrane. Because of the area scaling of microtubule number with cell size, this component must be inhibitory for microtubule nucleation, such that its sequestration leads effectively to an increased cytoplasmic concentration of nucleators with respect to volume scaling alone (Fig 4A). This type of model agrees quantitatively with our measurements of microtubule number in zebrafish spindles (Fig 4C and Supplemental information).

The sequestration model can also explain the hierarchical scaling of spindles, namely the transition from constant to scaling microtubule dynamics. The upper limit in microtubule growth velocity v_p_ is likely related to a saturation of microtubule plus ends with MAPs that promote microtubule growth (e.g. XMAP215 or EB proteins) (*22, 38, 39*). The measured decrease in microtubule growth velocity as cells and spindles become smaller suggests that the number of MAPs per microtubule decreases below a critical value, such that microtubule ends are not saturated anymore. Assuming that the number of MAPs promoting microtubule growth (N_MAP_) scales with cell volume, our theory predicts that the number of MAPs per microtubule decreases monotonically with cell size, leading to a quantitative decrease in microtubule polymerization velocity consistent with our data (Fig 4D). In summary, our measurements in zebrafish spindles suggest a model in which microtubule nucleation and dynamics scale differently with the surface-to-volume ratio of the cell. This distinct geometric scaling leads to the hierarchy of regimes of spindle scaling.

### Spindle scaling in encapsulated *Xenopus* egg extract spindles confirms that microtubule nucleation governs spindle scaling

Our theory makes two key predictions if components that regulate microtubule nucleation are not sequestered in the cell membrane. First, microtubule number should scale linearly with the cell volume, as in the simplest limiting component model. Second, microtubule dynamics should remain constant during scaling. This is because in this situation both microtubule number and components that regulate microtubule dynamics would scale with cell volume, and thus their ratio would remain constant, Fig 4B.

To test these predictions we turned to *in vitro* experiments with biochemically identical *Xenopus* egg extract spindles encapsulated in droplets of different sizes (*10, 11*) (Fig 5A and Materials and Methods). In these cell-like compartments, proteins cannot partition to the boundary (*24*). Thus, according to our theory, spindles in these droplets should scale solely by microtubule nucleation, whereas microtubule dynamics should remain constant.

**Fig. 5:**
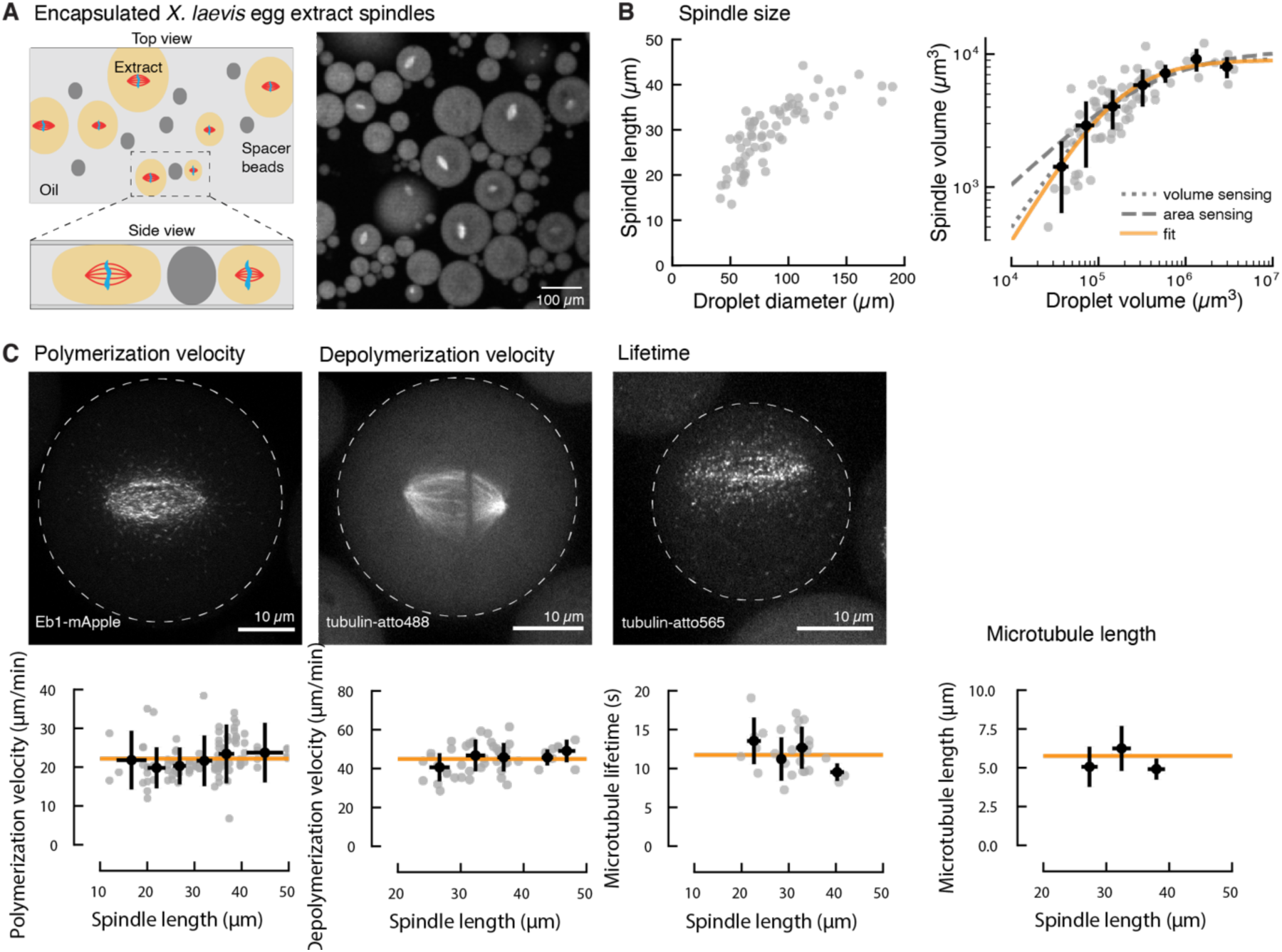
Spindle scaling in encapsulated *Xenopus* egg extract spindles. (A) Spindle assembly in cell-like compartments by encapsulating *Xenopus laevis* egg extract in inert oil. Droplets are compressed between spacer beads to improve image quality and spindle orientation. Microtubules are labeled with tubulin-atto488. (B) Spindle scaling with droplet size is consistent with a simple limiting component model in the absence of membrane partitioning. Left: Spindle length in encapsulated *Xenopus laevis* egg extract scales with droplet diameter. Each dot denotes an individual measurement (N=77). Right: Spindle volume scales linearly with droplet volume (gray dotted line and fit in orange). (C) Microtubule dynamics remain constant over a large range of spindle sizes in encapsulated *Xenopus laevis* egg extract., see also Movie S6-S8. Altogether these measurements determine the average microtubule length in the spindle (right). Gray dots denote individual measurements (N=32, 48, and 27), black dots are averages over 5 µm spindle size intervals. Error bars show standard deviations. The orange lines show the average values.

Consistent with our theoretical predictions, we found that the volume of encapsulated *Xenopus* egg extract spindles scales linearly with droplet volume, and not with surface area. Strikingly, in this assay microtubule dynamics are constant (v_p_ = 22 ± 6 µm/min, v_d_ = 45 ± 8 µm/min, and tau = 12 ± 3 s) irrespective of droplet size (Fig 5C-D and Movie S6-S8), in stark contrast to previous hypotheses of how these spindles scale (*10, 11, 22, 24*). Taken together, these results show that encapsulated *Xenopus* egg extract spindles scale exclusively by modulating microtubule number, and not by microtubule dynamics. Moreover, constant microtubule dynamics in the absence of boundary absorption suggests that membrane sequestration is indeed necessary to account for the hierarchical scaling of spindles.

## Discussion

Our quantitative measurements of microtubule dynamics, organization and nucleation in early zebrafish embryos and encapsulated *Xenopus* egg extracts demonstrate that spindle scaling is governed by microtubule nucleation. In contrast, microtubule polymerization dynamics only contributes as a secondary mechanism below a critical cell size (Fig 6). Importantly, even in small cells, changes in microtubule dynamics are insufficient to account for spindle scaling. Our results explain previous measurements in different organisms that appear contradictory within a unified framework. In the large cells of *Xenopus laevis* and zebrafish embryos, spindle size reaches an upper limit because of the diffusion-limited activation of microtubule nucleators (4), (*37*), (*15*). In intermediate-sized cells, spindles scale by limiting microtubule nucleation. Finally, in small cells microtubule dynamics also begin to contribute to spindle scaling (*17*).

**Fig. 6:**
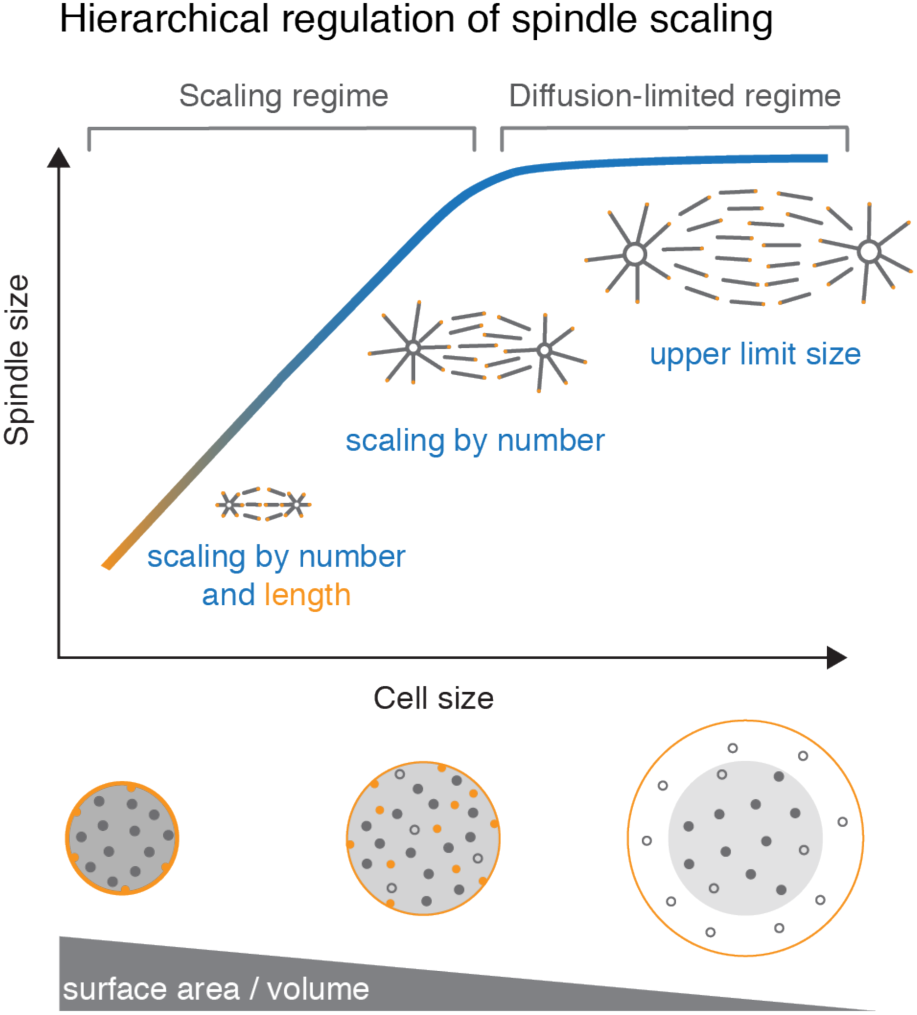
Hierarchical regulation of spindle scaling. Schematics of the different regimes of spindle scaling. For large cells, spindles reach an upper limit because activation and availability of microtubule nucleators is restricted to a reaction-diffusion volume. Below this volume, spindles scale with cell size by regulating microtubule number, which depends on the surface area of the cell through the sequestration of a microtubule nucleation inhibitor. When the surface to volume ratio exceeds a critical value, components that regulate microtubule dynamics become limiting, leading to a change in microtubule length for small cells.

Our data support a model in which the number of microtubules within the spindle is sensitive to the cell surface through an interplay of component limitation and sequestration of an inhibitor of microtubule nucleation to the cell boundary. This model quantitatively explains both the exact scaling of spindles with cell size and the hierarchy of nucleation over microtubule dynamics, and it is consistent with recent studies showing that importin alpha is sequestered in the membrane (*8, 24*). Importin alpha forms a complex with spindle assembly factors, and thus may effectively act as an inhibitor of nucleating factors that need to be released from importin by RanGTP (*40*-*42*). However, sequestration of other factors may contribute to the scaling we observe. For example, our data would also be consistent with a model where, apart from a nucleation inhibitor, MAPs are further limited by sequestration in the membrane (*8, 24*). This model would only work, however, provided that the sequestration effect contributes to a super-volume scaling (i.e., by sequestering components that promote microtubule growth, or inhibit depolymerizing factors). Importantly, our work restricts the class of molecular components that drive spindle scaling to those that regulate microtubule nucleation.

Our work shows that changes in microtubule dynamics are dispensable for spindle scaling. However, variability in spindle size across species for a given cell size could still be explained by differences in the overall microtubule dynamics (*7, 16, 17, 22*). We propose that these differences in microtubule dynamics, and possibly nucleation, reflect interspecies differences in relative protein amounts within a cell (*7, 43, 44*). These relative amounts in turn may set reference spindle sizes and microtubule lengths for each species—consistent with the organism-specific upper limit in spindle size (*12*)—but may not reflect the mechanisms that scale those spindles within the same species. This is because we found that spindle scaling is purely determined by changes in cell geometry, i.e. cell volume and surface area, that mainly affect microtubule number. Ultimately, this may explain why the shape of the scaling curves of spindles appears to be universal across species (*12*).

The hierarchical regulation of spindle size may help fulfil the architectural requirements of correct spindle function during the dramatic scaling that occurs in early embryo development (*45*). Scaling by nucleation ensures a constant microtubule density near chromosomes regardless of spindle size. In contrast, scaling by microtubule length would impose strong geometric constraints on the architecture of spindles. In large spindles, the microtubule density near the chromosomes would decrease dramatically with respect to small spindles, which may affect their mechanical integrity (*45*). Additionally, large spindles would require equally long microtubules, and thus increased stability that may be detrimental to the robustness and error correction in spindles (*46, 47*). However, scaling solely by microtubule nucleation sets a lower limit to spindle size, given by the length of microtubules. To keep scaling beyond this limit, microtubule length must decrease, which may explain the onset of the scaling of microtubule dynamics we observe for small cells. We thus speculate that hierarchical scaling allows for the proportionate change of microtubule architecture while maintaining the proper microtubule density throughout the structure, irrespective of spindle size. This mechanism does not require complex additional regulatory pathways and illustrates how cell geometry can instruct organelle size control.

## Acknowledgements

We acknowledge Martin Loose, Pavel Tomancak, Iain Patten, Nadine Vastenhouw, Christoph Zechner, Stephan Grill, and Otger Campàs for fruitful discussions and careful revision of the manuscript. We thank Maria Elsner, the Norden lab, the Vastenhouw lab, and Daniele Soroldoni for helping us with fish handling. We thank Heino Andreas for frog maintenance and the Light Microscopy Facility (LMF) and Fish Facility at MPI-CBG. This work was supported by the Human Frontiers Science Program (CDA00074/2014, to JB). The authors declare no competing interests.

## Supplementary Materials

Materials and Methods

Figs. S1 to S6

Movie Captions S1 to S8

Supplemental information

## Materials and Methods

### Animals

Zebrafish and *Xenopus laevis* adults and embryos were handled according to established protocols (*48, 49*). Zebrafish transgenic lines used in this study are Tg(bactin:GFP-utrophin) (*50*) and Tg(bactin:GFP-DCX) (*51*). The experiments were approved and licensed by the local animal ethics committee (Landesdirektion Sachsen, Germany; license no. DD24.1-5131/ 394/ 33 and DD24-5131/367/9) and carried out in accordance with the European Communities Council Directive 2010/63/EU on the protection of animals used for scientific purposes, as well as the German Animal Welfare Act.

### Zebrafish sample preparation

Zebrafish were maintained and bred at 27°C. Embryos were collected from pairwise mating in E3 medium (5mM NaCl, 0.17 mM KCl, 0.33 mM CaCl_2_, 0.33 mM MgSO_4_) within 15 min after spawning and kept at 24 – 28 °C. Embryo clutch quality was inspected on a dissection stereomicroscope and staged according to morphological criteria (*52*).

Fluorescently labeled proteins were injected into the yolk or cell of one-cell stage wildtype AB or transgenic zebrafish embryos according to (*53*). Injection volumes were carefully calibrated and were typically 1 – 1.5 nl. Protein concentrations were optimized based on dilution series (see sections below).

Pig tubulin was purified from pig brains following cycles of polymerization and depolymerization according to (*54*). Tubulin was labeled with fluorescent dyes (NHS-ester atto488 or atto565 (ATTO-TEC)) according to (*55*). Frog tubulin was purified from *Xenopus laevis* egg extract and directly labeled with NHS-ester atto565 (ATTO-TEC) according to (*56*).

### Quantification of spindle and cell volumes during early zebrafish development

To analyze cellular and spindle volumes, transgenic zebrafish embryos ubiquitously expressing a cell boundary marker (GFP-tagged actin-binding domain of human utrophin Tg(bactin:GFP-utrophin) (*50*)) were microinjected with fluorescently labeled pig tubulin-atto565 (concentration prior to injection: 15 µM, 90% labeled). For light sheet microscopy, embryos were manually dechorionated in E3 medium using watchmaker forceps (Dumont No.55) and mounted in 0.7 % low melting-point agarose (Sigma-Aldrich) in glass capillaries (Brand Transferpettor caps, inner diameter 1 mm). During image acquisition, the agarose column including the embryos was extruded from the glass capillary. Live imaging was performed on a Zeiss Light sheet Z.1 microscope, equipped with a 20x water-dipping objective (Zeiss Plan-Apochromat 20x, WI, NA 1.0) and a pco.edge sCMOS camera. Experiments were performed at 27.5 °C. Z-stacks spanning almost the entire embryo volume were acquired with 2 µm optical sectioning every 2 min throughout the first five hours of embryonic development (typically starting at 2 - 4 cell stage until epiboly onset).

The acquired dual color time-lapse movies of fluorescently labeled microtubules and cell boundaries were used to extract cell and spindle volume at different developmental stages. The centrosome positions (region of highest fluorescence at spindle poles) of metaphase spindles were manually annotated in Fiji (*57*). These positions were used as input for the custom-written Python scripts to automatically segment metaphase spindles and their corresponding cell.

For spindle segmentation, the image stack with fluorescently labeled microtubules was cropped around every annotated spindle. To aid visualization and subsequent data analysis, every spindle data set was rotated such that both spindle poles are horizontally aligned in the same image plane. Crucial for the following segmentation step was a background subtraction based on filtering local maxima with the scikit-image Python toolbox (*58*). Spindles were segmented in 3D using a histogram-based threshold (Otsu’s method). Segmented objects touching the boundary or below a threshold size were omitted. The segmentation quality was manually inspected. 3D spindle segmentation allowed extraction of the total spindle volume. To separate the total spindle volume into the spindle body and two spindle asters, the tubulin fluorescence intensity along a line connecting the centrosomes was measured and the local intensity minima between centrosomes and spindle body were automatically detected. The distance between these minima determined the pole-to-pole spindle length. Additionally, the average fluorescence intensity in the segmented spindle was calculated for 3 embryo data sets that were injected with a lower concentration of tubulin-atto565 (7 µM) and acquired at reduced laser power with fixed imaging parameters.

For cell segmentation, the Fiji plugin *LimeSeg* was used (*59*). In brief, at every annotated spindle a sphere (typically with a diameter equal to spindle length) was initialized as a seed for segmentation. These spheres expand depending on the user-defined ‘pressure’ acting on the surface (typical value 0.02) and the defined smallest feature size that *LimeSeg* detects (typically 2 – 3 pixel). Cell segmentation results were visually inspected. Finally, based on the centroid coordinates of the segmented objects every segmented cell was matched to its corresponding spindle and the scaling relationship between cell volume and spindle volume obtained.

### High-resolution imaging of microtubule dynamics

To study microtubule polymerization dynamics, wildtype AB zebrafish embryos were microinjected at the one-cell stage with EB1-GFP (concentration prior to injection: 20 µM). Embryos were mechanically dechorionated and mounted in 1% low melting-point agarose in E3 medium supplemented with 20% w/v Iodixanol (Sigma-Aldrich) to minimize refractive index mismatch (*60*) inside a 35 mm glass bottom petri dish (MatTek). The agarose patch was covered with E3 medium to prevent desiccation. Growing microtubule plus ends in the metaphase spindle were imaged using a spinning disk confocal microscope (Nikon Ti Eclipse, Yokogawa CSU-X1), equipped with a back-illuminated EMCCD camera (iXon DU-888 or DU-897, Andor) and a 60x water-immersion objective (CFI Plan Apochromat 60x WI, NA 1.2, Nikon). Image acquisition was controlled by Andor iQ software. Temperature was kept at 26.5 - 27.5 °C. Single z-planes of the spindle midplane were acquired every 0.5 – 2 s for a total of 0.5 – 2 min.

### Quantification of microtubule growth velocity

Microtubule growth velocity was inferred from kymographs of EB comets. Kymographs were drawn along the spindle long axis (centrosome to centrosome). Spindles that underwent significant sideways movement were excluded from analysis. Kymographs were analyzed either by manually tracing the lines of EB comets or by using the Fiji plugin *Directionality*. This plugin calculates the orientation of the input image based on Fourier component analysis and generates directionality histograms. These angle distributions were further processed in Python. Microtubule polymerization velocity was obtained by fitting Gaussian functions to the two peaks in the angle histogram (corresponding to the two populations of growing microtubule plus ends) and calculating the average velocity from that angle. Polymerization velocities were calculated individually for every analyzed spindle and averaged across 5 µm spindle length intervals.

### Tubulin speckle microscopy

To study microtubule turnover, transgenic zebrafish embryos expressing the GFP-tagged microtubule-binding protein doublecortin (Tg(bactin:GFP-DCX) were microinjected at the one-cell stage with frog tubulin-atto565 (concentration prior to injection: 1.6 µM, 22% labeled). The tubulin concentration for sparse tubulin labeling was optimized by dilution series. Embryos were mechanically dechorionated and mounted in 0.7% low melting-point agarose (Sigma) in glass capillaries (Brand Transferpettor caps, inner diameter 1 mm). Light-sheet microscopy was employed for *in vivo* imaging of tubulin speckles. During image acquisition, the agarose column including the embryos was extruded from the glass capillary. Imaging was performed on a Zeiss Light sheet Z.1 microscope with a Zeiss Plan-Apochromat 20x water-dipping objective (NA 1.0) at 27.5 °C. The doublecortin-GFP channel (Tg(bactin:GFP-DCX) was used to identify metaphase spindles. For tubulin speckle imaging, 5 z-planes around the spindle midplane were recorded with 1 µm spacing every 1.5 – 2.5 s for 0.5 – 2 min. Before and after speckle image acquisition (frog tubulin-atto565), a snapshot of the fully labeled spindle (Tg(bactin:GFP-DCX) was acquired to determine spindle length.

### Quantification of microtubule lifetimes

Single-particle tracking of tubulin speckles was performed using the tracking software *TrackMate* (*61*). Before tracking, metaphase spindles were manually cropped and a photobleaching correction was applied to the acquired time-lapse movies using Fiji. Depending on the particle density, single planes or maximum intensity projections of up to 5 neighboring z-planes were used for analysis. Spindles were segmented in Fiji by blurring the maximum intensity projections over time with a 5 - 10 pixel Gaussian filter, and applying a histogram-based threshold (Mean or <u>IsoData</u>). Alternatively, regions of interest were selected manually in Fiji. *TrackMate* detects particles based on DoG (Difference of Gaussian) segmentation. Particle detection was visually inspected and parameters for the blob diameter (typically 2 pixel) and threshold were refined, if necessary. Then sequences of single, localized events were linked to trajectories based on the assumption of linear motion (linear motion LAP tracker). Typical values were 2 pixels for the initial search radius, 2 pixels for the search radius and 1 frame maximum frame gap. The tracks were exported as .xml file and additional analysis and data fitting was performed using custom-written MATLAB code. Only tracks that appeared and disappeared during the length of the acquired time-lapse and that existed for at least 2 frames were included in the analysis. To obtain the speckle lifetime distribution, the tracks were binned with a bin size equal to the frame rate (typically 2 s). The speckle lifetime distributions P(t) per spindle size interval were fit to a diffusion with drift process *P(t)* = *a t*^-3/2^*e*^*-t*/4*τ*^, where *τ* is the lifetime of a microtubule of average length (*30, 62*). In practice, we fit the log of the relative frequencies. In total, 47 spindles were analyzed. We estimated the change in lifetime due to the observed changes in polymerization velocity using the formula 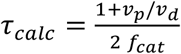, where *f*_*cat*_ is the catastrophe frequency (solid line in Fig. 3C). We calculated the catastrophe frequency from an exemplary spindle of length L_spd_ = 30 − 35 µm, that yield a catastrophe rate of 0.16 s^−1^. We used the measured microtubule dynamics (*v*_*p*_, *v*_*d*_, *τ*) to estimate the average microtubule length L_MT_ in the spindle according to 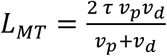

### *L*aser-ablation induced microtubule depolymerization

To locally induce microtubule depolymerization, wildtype AB zebrafish embryos microinjected with pig tubulin-atto565 were mechanically dechorionated and mounted in 1% low melting-point agarose (Sigma) in E3 medium supplemented with 20% w/v Iodixanol (Sigma) for refractive index matching inside a glass bottom petri dish (MatTek). The agarose patch was covered with E3 medium to prevent desiccation. Fully labeled metaphase spindles were imaged using a spinning disk confocal microscope (Nikon Ti Eclipse, Yokogawa CSU-X1) equipped with an EMCCD camera (iXon DU-888 or DU-897, Andor) and a 60x/1.2 NA water immersion objective. Image acquisition was controlled by Andor iQ software.

For laser ablation, a mode-locked Ti:Sapphire laser (Coherent Chameleon Vision II) was coupled into the back port of the Nikon spinning disk microscope. The custom-built laser ablation setup is based on a previously described layout (*63*). The Ti:Sapphire laser delivers 140 fs pulses at a repetition rate of 80 MHz. The output laser power was manually modulated using a half-wave plate and a Glan-Thompson polarizer. Additionally, an acousto-optical pulse picker (pulseSelect, APE) was added the optical path to reduce the repetition rate to 20 kHz, which reduced photodamage significantly. Laser ablation was performed using a wavelength of 800 nm and typically a power of 350 µW after the pulse picker. Line cuts perpendicular to the spindle long axis were implemented by moving the sample with a high-precision piezo stage (PInano) relative to the stationary cutting laser. The cutting procedure was automatically executed by a custom-written software that controlled the piezo-stage and a mechanical shutter in the optical path. Several line cuts were performed in depth around the focal plane to enhance ablation efficiency. The length and depth of the cut were adjusted relative to spindle size (typically, length 10 µm and depth 2 µm). Every metaphase spindle was cut only once. To capture the fast dynamics of microtubule depolymerization, single z-planes of the spindle midplane were acquired every 0.2 – 0.5 s for a total of 30 – 60 s following ablation. Spindles recovered the microtubule network at the ablation site within 30 s. The subsequent cell divisions proceeded normally after ablation.

### Laser ablation analysis

Microtubule depolymerization following femtosecond laser ablation was analyzed using a custom-written MATLAB code. The total amount of microtubules that depolymerized during the time interval δt was calculated by subtracting raw images of the acquired time-lapse movie with a time difference δt of 2 – 3 s from each other and integrating these differential intensities perpendicular to the spindle long axis. Depending on the position of the cut, the integrated differential intensities showed one or two well-defined peaks (due to microtubule polarity at the cut site). Each peak was fit to a Gaussian function, 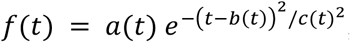, to quantify the position of the maximum b and the area, 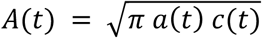, under the curve, where a is the amplitude and c is the width of the Gaussian at time t after the cut. Microtubule polarity p at the position of the cut was measured by calculating the relative ratio of the area under the two integrated differential intensity peaks immediately after the cut, *p* = *A*_1_ / (*A*_1_ + *A*_2_), where *A*_1_and *A*_2_ are the areas under the two depolymerization fronts at t = 0. If only one depolymerization front was detected, the other one was set to zero. The polarity profiles were fit to a logistic function, *p* = 1/(1 + *e*^−*a*(*x*−*b*^)), where *a* is the steepness and *b* the midpoint of the curve. The steepness *a* was used to infer the width, *w* = 2 *a*^−1^, of the overlap region of microtubules of opposed polarity (Fig S3). Following ablation, the peaks moved toward the nearest pole. The position of the maxima of the Gaussian fits over time was fit to a line to infer the microtubule depolymerization velocity *v*_*d*_. The area under the Gaussian is proportional to the total amount of depolymerized microtubules of the same polarity during the time interval δt. The initial increase in area due to the bleach mark of the laser cut was ignored for the subsequent analysis (∼2s after cut). The areas were normalized to the maximum value. The area decreased monotonically over time as the depolymerization front moved away from the cut and was fit by an exponential *A*(*t*) = *A*(0)*e*^−*t*/*v*^. To compare the exponential decay time *v* for different spindle sizes, individual measurements of *v* were averaged across spindles within a length interval of 5 µm. The microtubule depolymerization velocity *v*_*d*_ was used to express the exponential decay time *v* as a function of the distance from the cut (exponential decay length *λ* = *v v*_*d*_). The decay length *λ* at the position of the cut is proportional to the number of minus ends per unit length at that location. Finally, the relative microtubule number for differently sized spindles was obtained by multiplying the averaged decay length *λ* with the average spindle mass M_spd_ for every spindle size bin. In total, 162 cuts were analyzed in spindles ranging from 15 – 55 µm in aster-aster length, but only area decays with an R^2^ > 0.8 of the exponential fit were considered for further analysis. This restriction reduced the number of analyzed data sets to 98.

### Microtubule density profiles

To obtain the fluorescence intensity profiles along the spindle long axis, spinning disk images of tubulin-labeled spindles (pig tubulin-atto565) that were acquired prior to laser ablation were used. Centrosomes were annotated manually in Fiji. Image analysis was performed in MATLAB. To correct uneven background illumination, a top-hat filter with a disk-shaped structural element (100 pixel) was applied to the image. Following background subtraction, the fluorescence intensity perpendicular to the spindle long axis was integrated to yield the spatial intensity profile along the spindle long axis. The integrated fluorescence intensity along the spindle long axis was divided by the spindle width at every position, which was obtained by segmenting the spindle mid-plane (Otsu thresholding). The fluorescence intensity profiles were normalized and several profiles of similar aster-aster spindle length were aligned to the spindle center and averaged together. The amplitudes of the normalized microtubule density profiles were rescaled by the total average fluorescence intensity in the spindle (as measured from SPIM data).

To characterize the size of spindle asters, the radial intensity profiles around the centrosomes were calculated. The radial fluorescence intensity profile of the spindle midplane was calculated in a half circle facing away from the spindle body, with the centrosome at the center. For each aster, the fluorescence intensity was radially averaged as a function of the distance from the centrosome. The averaged radial intensity profiles were fit to an exponential function to infer aster size from the exponential decay length.

### Quantification of microtubule transport

To quantify motor-driven microtubule transport in the spindle, a bleach mark was generated in fluorescently labeled zebrafish spindles (pig tubulin-atto565) and its motion towards the pole was analyzed. Experiments were carried out similar to previously described laser cuts, with the only difference that the output laser intensity of the cutting laser was reduced such that fluorescently labeled microtubules were bleached but no depolymerization was induced. The motion of the bleached line was quantified by generating kymographs along the spindle long axis and manually tracing the angle of the bleached region in Fiji. This angle was used to calculate the velocity of microtubule transport. In total, 27 bleaching experiments over a large range of spindle sizes were analyzed.

### Cytoplasmic extract preparation, *<u>in vitro</u>*<u>spindle assembly and encapsulation</u>

Cytostatic factor-arrested (CSF) cytoplasmic extracts were prepared from freshly laid eggs of *Xenopus laevis* and used for spindle assembly reactions as described in (*49*). Briefly, unfertilized *Xenopus laevis* oocytes arrested in metaphase of meiosis II were collected, dejellied and fractionated by centrifugation. The cytoplasmic layer was isolated and supplemented with protease inhibitors (LPC: Leupeptin, Pepstatin, Chymostatin) and Cytochalasin D (CyD) to a final concentration of 10 µg/ml each. Single reactions were cycled to interphase by adding frog sperm (300 - 1000 sperm/µl final concentration) and 0.4 mM Ca^2+^ solution. Reactions were incubated for 1 h. While fresh CSF extract containing LPC and CyD was kept on ice, all incubation steps were performed at 18 – 20 °C. Prior to initiation of spindle assembly, atto488 labeled purified pig tubulin was added to the reactions to a final concentration of 500 nM to visualize spindles. To visualize microtubule plus ends, EB1-mApple was added to a final concentration of 100 nM. For tubulin speckle microscopy, frog tubulin-atto565 was added to a final concentration of 100 nM (20% labeling ratio). Spindle assembly reactions were initiated with the addition of 1.3 volumes of fresh CSF extract (containing LPC and CyD) to labeled interphase extract. Spindle reactions were immediately encapsulated in droplets by slowly pipetting extract into Novec HFE-7500 oil supplemented with 2% (w/w) Pico-Surf surfactant (Sphere Fluidics) and flicking the reaction until the dispersion appears homogeneous and white. This procedure typically leads to extract droplets of 10 – 200 µm in diameter. Encapsulated spindle assembly reactions were incubated for 45 min at 18 – 20 °C prior to imaging.

### Image acquisition of encapsulated extract spindles

Spindle droplets were transferred by gentle pipetting to a microscope slide that was complemented with 45 µm spacer beads (Polybead Microspheres) to prevent squishing of droplets. The squash was covered with a Hellmanex-cleaned cover slip and sealed with Valap. Compression of spindle droplets led to a preferred orientation of the spindle long axis parallel to the imaging plane. Spindle droplets were imaged using a spinning disk confocal microscope (Nikon Ti Eclipse, Yokogawa CSU-X1), equipped with an EMCCD camera (iXon DU-897, Andor) and a 60x water-immersion objective at 20 °C. Image acquisition was controlled by Andor iQ software. For microtubule plus end tracking, single z-planes of EB1-mApple labelled spindles were acquired every 0.5 s. For tubulin speckle microscopy, single z-planes of frog tubulin-atto565 labeled spindles were imaged. To increase the signal-to-noise ratio of tubulin speckles, it was essential to average 4 consecutive 500 ms exposures. To measure spindle size, z-stacks of tubulin atto488-labeled spindles were acquired at a z-spacing of 1 – 2 µm. Image segmentation, laser ablation, and analysis to measure microtubule depolymerization, growth velocity, and lifetime was performed similar to zebrafish spindles.

**Fig. S1:**
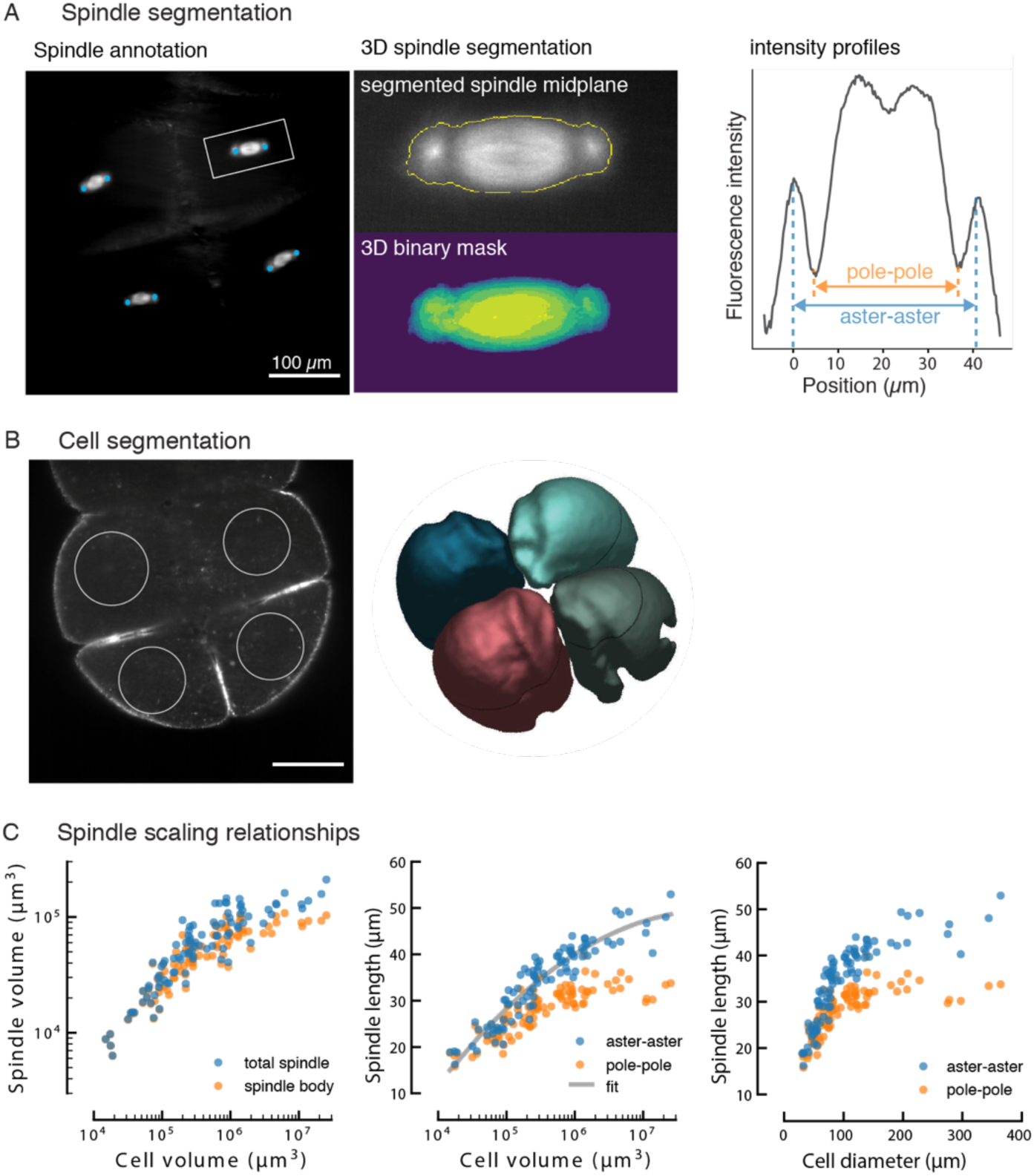
Analysis of spindle and cell sizes during early zebrafish development. (A) 3D segmentation of spindles during early zebrafish development. Left: The centrosome positions of metaphase spindles (tubulin-atto565) were manually annotated (blue dots). Middle: Every annotated spindle was segmented in 3D to determine the spindle volume. Exemplary image of the spindle midplane overlaid with contour of binary spindle mask (yellow line) and 3D representation of segmented spindle. Right: Fluorescence intensity profile along the spindle long axis: The annotated centrosome positions define the aster-aster spindle length, while the distance between the intensity minima between centrosomes and spindle body determine the pole-to-pole spindle length. (B) Cell segmentation. Left: Single plane of the fluorescently labeled cell boundary (utrophin-GFP). Scale bar, 100 µm. At every annotated spindle a sphere (white circle) was initialized as seed for the cell volume segmentation using LimeSeg. Right: Volume rendering of the segmented cells (see also Movie S2). (C) Spindle scaling during early zebrafish development: Spindle volume and length correlate with cell size in in small cells but reach an upper limit in larger cells. Blue dots denote the total spindle volume (or aster-aster length), orange dots denote the spindle body volume excluding asters (or pole-pole length). Cell diameter was calculated from the measured cell volume under the assumption that cells are spherical. To characterize the relationship between aster-aster spindle length L_spd_ and cell volume V, we fitted a 2^nd^ order polynomial function (*L*_*spd*_ = *a* log(*V*)^2^ + *b* log(*V*) + *c*, gray line) to the log-linear data. The fit parameters (a = −98.17, b = 15,83, c = −0,4217) were used to calculate the cell volume for experiments, where only spindle length was measured (i.e. microtubule dynamics and nucleation).

**Fig. S2:**
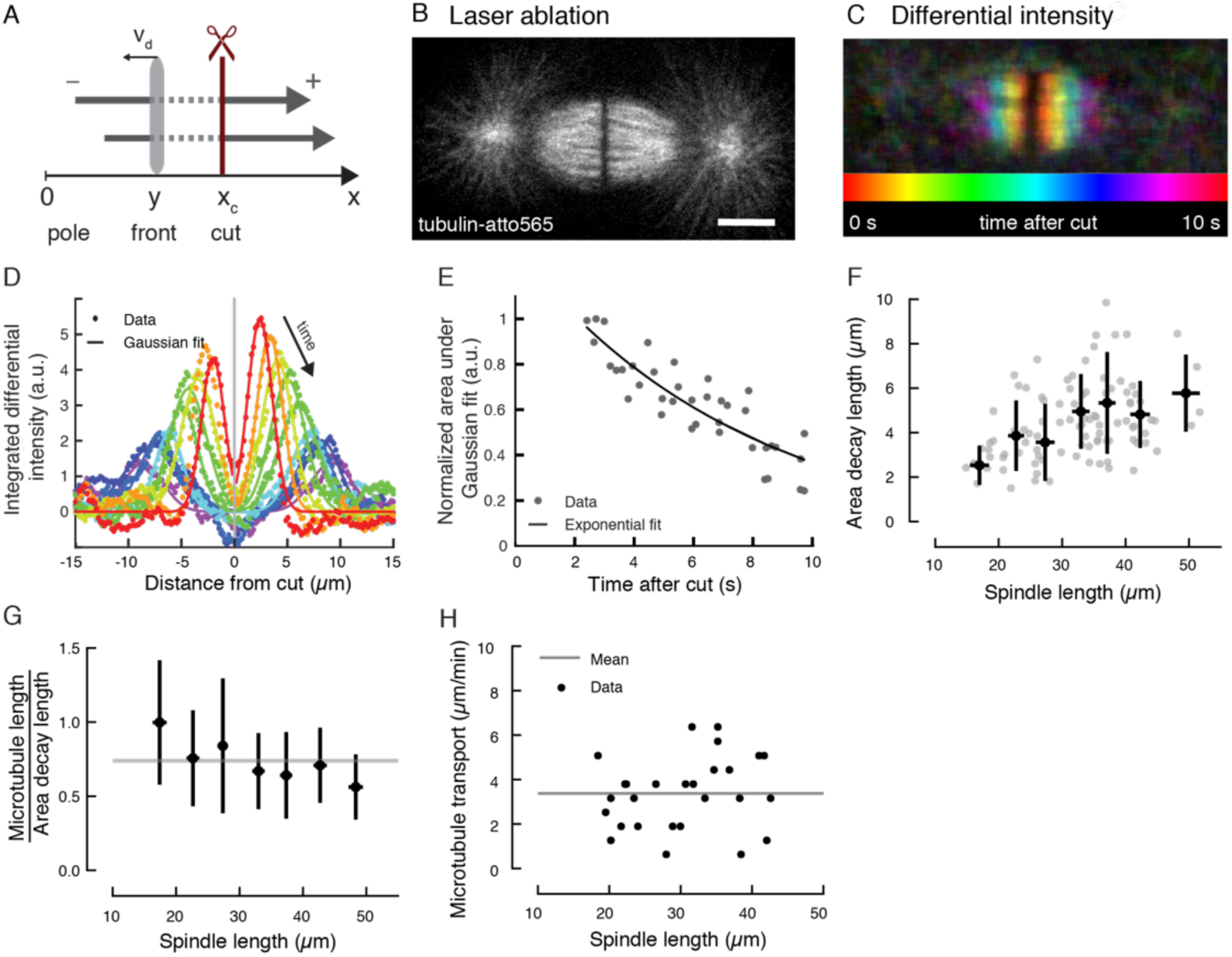
Analysis of laser-induced microtubule depolymerization. (A) Schematic of laser ablation. Severing microtubules (gray arrows) at position x_c_ induces synchronous depolymerization of the newly created plus ends. The depolymerization front (light gray area) moves at velocity v_d_ away from the cut site. (B) Linear laser cut in a spindle labeled with tubulin-atto565. Scale bar, 10 µm. (C) The differential fluorescence intensities reveal the extent of microtubule depolymerization at selected time points after ablation. Depolymerization fronts are color-coded by time. (D) The integrated differential intensities exhibit distinct peaks (color-coded by time) that move away from the cut. A Gaussian (solid line) was fit to each peak to quantify the area under these curves. (E) The area under the Gaussian fits is proportional to the amount of depolymerizing microtubules. Its exponential decrease following ablation indicates that microtubule minus ends are distributed throughout the spindle. (F) The area decay length (obtained from exponential fit in E) of the laser-induced depolymerization fronts is constant in spindles larger than 35 µm aster-aster length. In smaller spindles, the area decay length decreases with decreasing spindle size. Gray dots denote individual measurements. Averages over 5 µm spindle size intervals are show in black (mean ± SD). (G) The ratio between average microtubule length L_MT_ and the area decay length *λ*, which is proportional to the microtubule minus end density, remains constant during scaling. Thus, the change in microtubule length in small spindles is compensated by a change in microtubule number. The gray line shows the average of the ratio L_MT_ /*λ* over all spindle sizes. Error bars denote standard deviations. (H) Microtubule transport in differently sized spindles. Microtubules move on average at a velocity of 3.4 ± 1.6 µm/min due to motor-driven transport (gray line). Black dots denote individual measurements in spindles of different sizes.

**Fig. S3:**
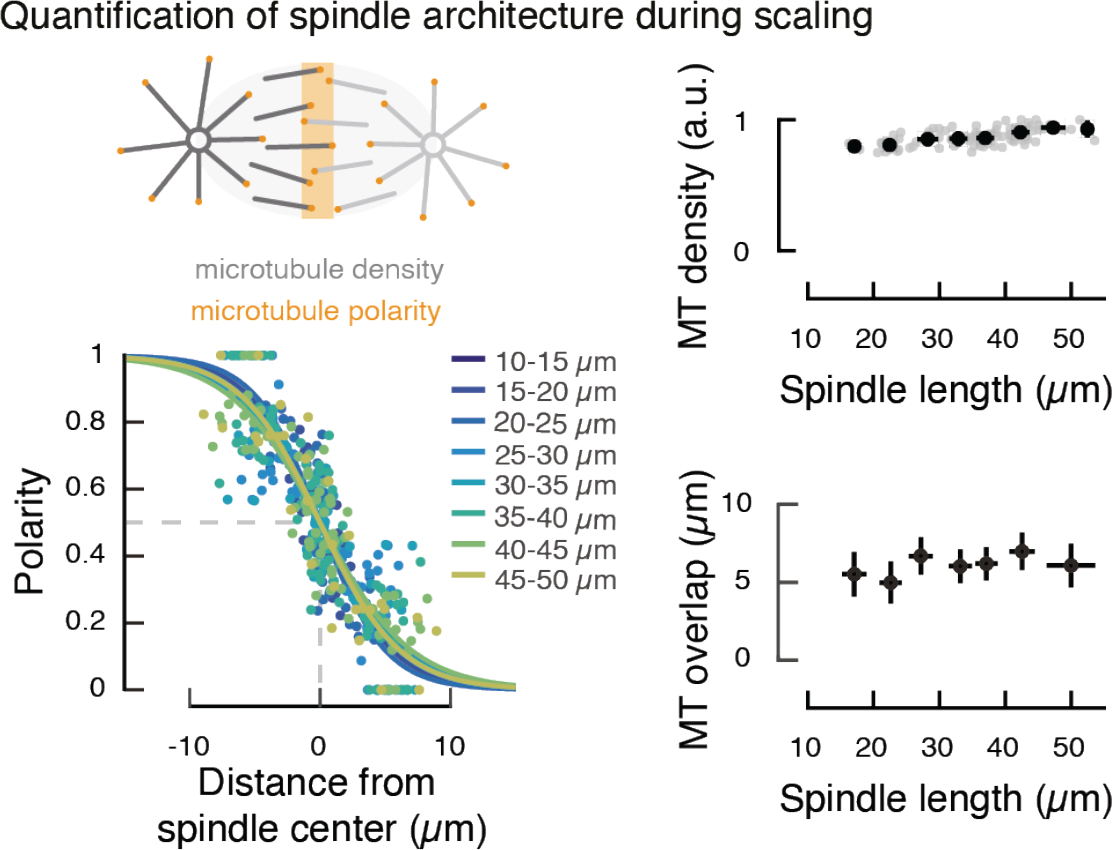
Spindle architecture remains unchanged during scaling. Upper left: The organization of microtubules within the spindle is characterized by the microtubule density and their polarity. The spindle is composed of two microtubule networks of opposed polarity (shown in light and dark gray) that may overlap at the spindle center (microtubule overlap highlighted in orange). Upper right: The average microtubule density (based on tubulin fluorescence intensity) decreases linearly with decreasing spindle size. Gray dots denote individual measurements (N = 96 in 3 different embryos) and black dots are averages over a 5 µm spindle size range. Error bars denote standard deviations. Lower left: Microtubule polarity profiles measured by laser ablation (see Supplemental Information) for spindles of different sizes. Each dot represents an individual cut at a certain distance from the spindle center (color-coded by spindle length, number of cuts N_cut_ = 190). At the spindle center, an equal number of microtubules points towards the chromosomes, whereas away from the spindle center all microtubules have the same polarity. The logistic fits (solid lines, color-coded by spindle length) characterize the overlap of microtubules of opposed polarity at the spindle center. Lower right: The region of microtubule overlap is narrow in comparison to spindle length and does not change during scaling.

**Fig. S4:**
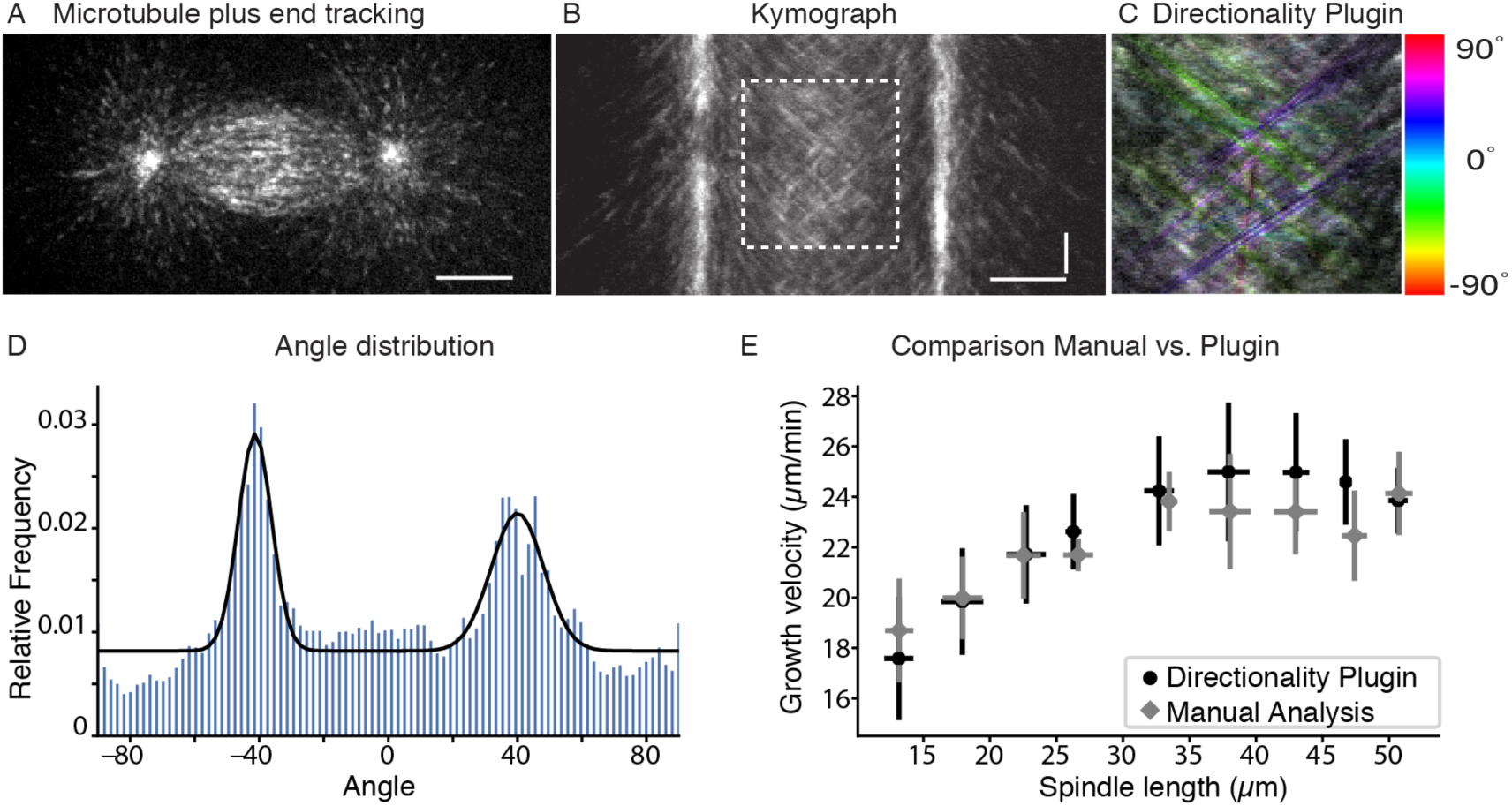
Quantification of microtubule growth velocities in metaphase spindles. (A) Exemplary metaphase spindle in the living zebrafish embryo fluorescently labeled with EB1-GFP to visualize growing microtubule plus ends. Scale bar, 10 µm. (B) Kymograph along a line connecting both centrosomes in the spindle. The horizontal axis represents distance and the vertical axis time (horizontal scale bar, 10 µm; vertical scale bar: 5 s). The bright vertical lines are the spindle poles. The motion of EB comets is apparent as lines at a certain angle both within the spindle body and away from the poles. (C) Manually specified regions within the kymograph (dotted rectangle in (B)) were analyzed using the Fiji Plugin Directionality. The colored lines represent the identified directions of the input image. (D) Histogram of the angle distribution obtained from (C). The two peaks are related to the two populations of EB comets moving from each pole towards the spindle center. Each peak was fit to a Gaussian function (black line) to infer the average orientation of the EB comets in the kymograph. (E) The microtubule growth velocity was calculated based on the inferred angles and is shown as a function of spindle length (aster-aster). The growth velocity obtained from the semi-automated analysis approach (black dots) agrees well with the manual kymograph analysis (gray diamonds). Error bars denote standard deviation.

**Fig. S5:**
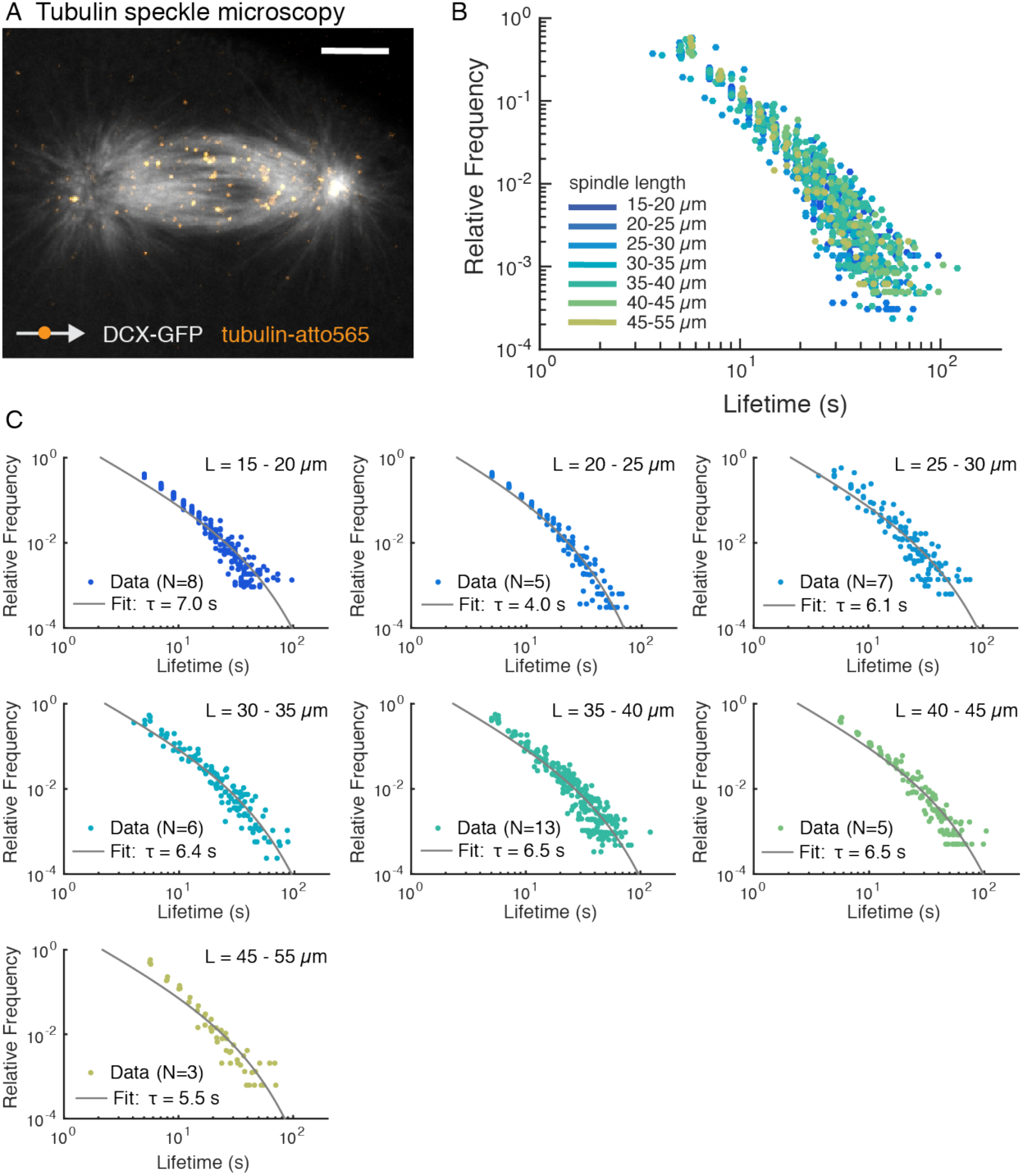
Quantification of microtubule lifetime distributions. (A) Exemplary snapshot of a fully labeled metaphase spindle (Tg(bactin:GFP-DCX), in gray) with sparsely labeled tubulin speckles (frog tubulin-atto565, in orange). Scale bar, 10 µm. (B) Speckle lifetime distributions for different spindle sizes. (C) The measured distributions for every spindle size *L* were combined and fit according to *P*(*t*) = *a t*^−3/2^*e*^−*t*/4*τ*^, where *τ* is the lifetime of a microtubule of average length (gray line). Microtubule lifetime *τ* is shown for every spindle size. *N* denotes the number of analyzed spindles.

**Fig. S6:**
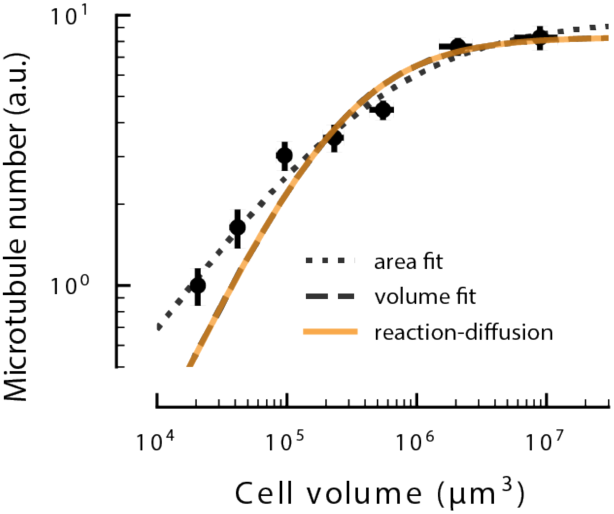
Comparison of the phenomenological volume fit with the full solution of the reaction diffusion problem in 3D for the simple limiting component. Fits of the full solution of the reaction diffusion of microtubule number (solid orange), and phenomenological volume and saturation fit (long dotted line). The phenomenological area fit (short dotted line) is shown for completeness.

## Supplementary Movies

**Movie S1 – Cell-size dependent spindle scaling during early zebrafish development.**

**Movie S2 – 3D view of spindles in a 8-stage embryo.**

**Movie S3 – Growing microtubule plus ends in a zebrafish spindle.**

**Movie S4– Laser ablation of a zebrafish spindle.**

**Movie S5 – Tubulin speckle microscopy in a zebrafish spindle.**

**Movie S6 - Growing microtubule plus ends in an encapsulated *X. laevis* egg extract spindle.**

**Movie S7 – Laser ablation in an encapsulated *X. laevis* egg extract spindle.**

**Movie S8 – Tubulin speckle microscopy in an encapsulated *X. laevis* egg extract spindle.**

## Supplemental Information SM1

### SM1. Laser ablation method

We used laser ablation to locally induce microtubule depolymerization. Analysis of the depolymerization wave allows quantification of the microtubule depolymerization velocity, microtubule polarity and the density of microtubule minus ends in the spindle (15, 27).Here, we describe the theoretical basis as well as the practical implementation of the method for the current work.

The laser ablation method relies on the observation that severing a microtubule by laser irradiation creates a new stable minus end and a new dynamic plus end. The newly created plus end immediately depolymerizes to the original minus end of the cut microtubule (Fig S2 A). In the spindle, laser ablation of fluorescently labeled microtubules induces synchronous microtubule depolymerization towards the nearest pole, which is visible as loss in fluorescence intensity (Fig S2 B-C). We quantified the extent of microtubule depolymerization by subtracting the raw fluorescence intensities of the acquired time-lapse images with a time difference *δt* from each other and integrating the differential intensities perpendicular to the spindle long axis. The area *A*(*t, x*_*c*_) under the integrated differential intensity peaks corresponds to microtubules of the same polarity that depolymerized during the time interval *δt*,

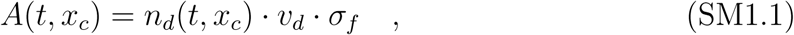

where *n*_*d*_ is the number density of depolymerizing microtubules from a cut at position *x*_*c*_, *v*_*d*_ is the depolymerization velocity and *σ*_*f*_ is the fluorescence per unit length and microtubule. We quantified the amount of microtubule depolymerization following ablation by fitting a Gaussian function to the peaks in the integrated differential intensities (Fig S2 D). The area *A*(*t, x*_*c*_) under the Gaussian, which is proportional to the amount of depolymerizing microtubules per unit time, decreases exponentially as the depolymerization front moves away from the cut, *A*(*t, x*_*c*_) = *A*(0, *x*_*c*_) · exp(−*t/v*), see Fig S2 E. The monotonic decrease indicates that progressively fewer microtubules depolymerize as the front advances. The number of depolymerizing microtubules *n*_*d*_ (per cross-section) changes, when depolymerization reaches a microtubule minus end,

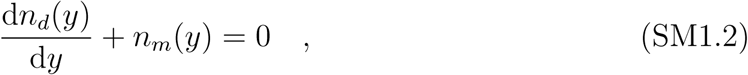

where *n*_*m*_*(y*) is the density of microtubule minus ends at a distance *y = v*_*d*_ · *t* from the cut. Thus, the change in the area *A(y, x*_*c*_) as the front of microtubule depolymerization moves away from the cut contains valuable information on the density of microtubule minus ends *n*_*m*_ in the spindle

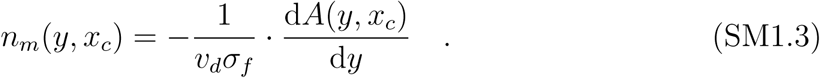

By taking the exponential decay of *A*(*y, x*_*c*_) = *A*(0, *x*_*c*_)·exp(-*y*/,*λ*) with a characteristic decay length *λ* and equation SM1.1 into account, the minus end density *n*_*m*_*(x*_*c*_) at the cut position (*y* = 0) is then given by

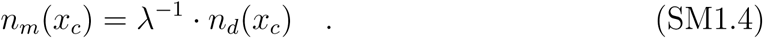

Thus, the density of microtubule minus ends at the cut site *n*_*m*_*(x*_*c*_) is proportional to the inverse of the decay length, *λ* and the number of depolymerizing microtubules *n*_*d*_(*x*_*c*_) (per cross-section) at the cut site. We assume that the number density of depolymerizing microtubules *n*_*d*_ is proportional to the average microtubule density *ρ* in the spindle.

Finally, the number of microtubule minus ends *N*_MT_ in the spindle can be obtained by integrating the minus end density *n*_*m*_(*x*_*c*_) over the spindle volume *V*_spd_

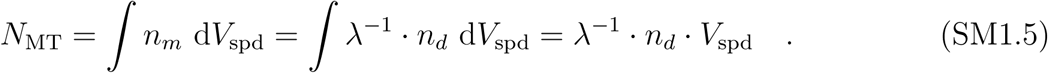

Here, we used the fact that in our measurements, *λ* is constant within a spindle (see Fig S4), and *ρ* is also constant (based on our measurements of microtubule density using tubulin fluorescence intensity). This implies that the microtubule minus end density *n*_*m*_ across the spindle is constant. In that case, microtubule number *N*_MT_ is given by *N*_MT_ = *λ*^−1^ · *ρ* · *V*_*spd*_. This implies that the number of microtubules *N*_MT_ in the spindle is proportional to spindle mass, *M*_*spd*_ = *ρ* · *V*_spd_, and the exponential decay length *λ* of the laser-induced microtubule depolymerization

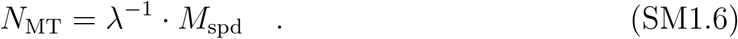

To quantify microtubule number *N*_MT_ during scaling by laser ablation, we performed laser cuts in differently sized spindles during early zebrafish development and analyzed the laser-induced microtubule depolymerization fronts as described above. These experiments revealed that the area decay length *λ*, which is proportional to the microtubule minus end density at the cut site decreases with decreasing spindle size (Fig S2 F). In spindles larger than 35 µm aster-aster length (regime of constant microtubule length), the area decay length *λ* also plateaus. Importantly, SM1.6 implies that, during spindle scaling, the change in the area decay length *λ* has to compensate the change in average microtubule length *L*_MT_. This is because spindle mass is determined by the length and number of microtubules, *M*_*spd*_ ∼ · *N*_MT_ · *L*_MT_, and therefore, *L*_MT_/*λ* = 1. Indeed, we found that the ratio between average microtubule length and area decay length is relatively constant during scaling (see Fig S2 G). The length scale *λ*, at which the microtubule depolymerization front decays whenever it reaches a microtubule minus end, is comparable to the measured average microtubule length. The small difference between *L*_MT_ and *λ* may be because of an underestimation of microtubule length based on the dynamics measurements. Nevertheless, the change in microtubule length during scaling is compensated by a change in microtubule number.

### Spatial distribution of microtubule nucleation

Here, we investigated the spatial distribution of microtubule nucleation in the spindle. Laser cuts performed at different locations within the spindle reveal the local microtubule minus end density (see above). In the case of negligible transport, the microtubule minus end density is a reasonable approximation of microtubule nucleation sites. We quantified motor-driven microtubule transport in the spindle by generating a bleach mark in fluorescently labeled zebrafish spindles and quantifying the motion of this bleach mark. We found that microtubule transport occurs at 3.4 *±* 1.6 µm*/*min in zebrafish spindles and is independent of spindle size, see Fig S2-H. This rate is comparably slow to the short microtubule lifetime of ∼5 s, such that a microtubule will move less than 0.5 µm away from its nucleation site. Consequently, the location of microtubule minus ends corresponds to microtubule nucleation sites.

### Microtubule minus end density

The microtubule minus end density *n*_*m*_ of microtubules with the same polarity is proportional to the change in the total amount of microtubule depolymerization, which is characterized by the exponential area decay length *λ*, see equation (SM1.4). We performed laser cuts at different locations within the spindle and found that the area decay length *λ* is independent of the cut position. The constant area decay length *λ* indicates that microtubule minus end density *n*_*m*_ is constant throughout the spindle body. However, not all locations within the spindle were accessible by laser ablation. We could reliably analyze laser cuts near the spindle center. Near the spindle poles, however, the laser-induced depolymerization fronts disappeared too quickly due to the fast depolymerization velocity to infer the area decay. Therefore, the relative microtubule minus end density could only be inferred by laser ablation near the spindle center. In this region, the microtubule minus end density is constant.

### Microtubule density

The constant microtubule minus end density in the spindle body (measured by laser ablation) is consistent with the tubulin fluorescence intensity profiles, which we assumed to be proportional to microtubule density. Because microtubule transport is negligible and microtubules are short relative to spindle size, these microtubule density profiles are a close approximation of the nucleation profiles in spindles.

### Microtubule polarity

To infer the nucleation profile of microtubules of a given polarity, i.e. of all microtubules with minus ends pointing in the same direction, we investigated the spatial organization of microtubules in the spindle. Laser ablation reveals the polarity of microtubules in the spindle body, because the total amount of microtubule depolymerization immediately after the cut, *A*(*t* = 0), is typically different for the laser-induced depolymerization fronts. This asymmetry is caused by a different number of microtubules of opposed polarity at the cut site. The relative fraction of the integrated intensity loss for the two depolyermization fronts immediately after the cut reveals microtubule polarity at the location of the cut. At the spindle center, microtubule polarity is 0.5, with equal numbers of microtubules pointing in both directions, whereas close to a pole, the majority of microtubules are oriented with their plus ends away from the pole. We fitted a logistic function to the polarity profile to characterize the width of the overlap region of microtubules with opposed polarity.

### Microtubule nucleation

Finally, we inferred the nucleation profile of microtubules of the same polarity by multiplying the microtubule density profile with the polarity profile. To estimate a characteristic length scale of microtubule nucleation in the spindle body, we fitted a Gaussian function to the nucleation profiles (excluding centrosomes). The extent of microtubule nucleation in the spindle body increases linearly with increasing spindle size. This increase is comparable to the increase in aster size, see Fig 3.

## SM2. Theory of scaling of microtubule number and dynamics with cell volume

Here, we propose a model that captures the hierarchical regulation of spindle size by microtubule nucleation and dynamics. Specifically, our model recapitulates (1) the scaling of microtubule number with cell size and (2) the transition from constant to scaling microtubule dynamics below a critical cell size. We first consider the reaction-diffusion of a simple limit component and then include the effect of the sequestration of a nucleation inhibitor in the cell boundary.

### Reaction-diffusion of a simple limiting component mechanism for microtubule nucleation

To characterize the scaling of microtubule number with cell volume, we consider a mechanism in which microtubule number is proportional to the number of active nucleators in the cell. Nucleators are locally activated near chromosomes through the Ran pathway and diffuse through the cytoplasm, where they can become inactivated or bind to other microtubules to nucleate other microtubules (14, 15, 33, 37).Local activation, diffusion and degradation set up a gradient with a characteristic length scale that is equal to the square root of the ratio of diffusion *D* to degradation rate 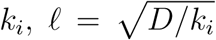. For simplicity, we ignore any coupling between microtubule nucleation and dynamics (14, 15, 33) which may alter the gradient of active nucleators. In our theory, the amount of microtubules will be proportional to the amount of active nucleators, with a proportionality constant that reflects this detailed interaction and kinetics of nucleators and microtubules. Because our data shows that microtubule dynamics are mainly invariant with cell size and only exhibit a subtle change below a critical cell size, the main driver for changing microtubule number arises from a limiting amount of nucleators and not because of changes in microtubule dynamics—that in turn affect nucleation. Therefore, here we consider only the population of active nucleators and assume microtubules are proportional to them with a constant of proportionality that does not depend on cell size.

In large cells, the length scale of the reaction-diffusion limited zone of active nucleators may be smaller than the cell diameter. In this regime, the availability of nucleators does not depend on cytoplasmic volume. Consequently, the amount of active nucleators will remain constant as long as cell volume *V*_*cell*_ is larger than the reaction-diffusion volume *V*_0_. In cells smaller than *V*_0_, activation of nucleators is not limited by diffusion. In this regime, components are well-mixed and will decrease with decreasing cell volume. Thus, this type of model can conceptually capture the saturation regime of spindle size and the transition to a scaling regime. The corresponding reaction-diffusion process in spherical coordinates in three dimensions reads,

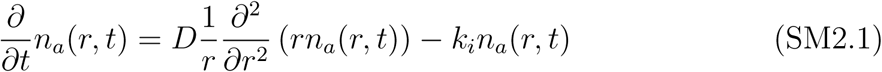

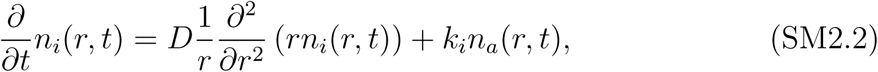

where *n*_*a*_ and *n*_*i*_ are the concentrations of active and inactive nucleators, respectively, *D* is the diffusion constant of active and inactive nucleators and *k*_*i*_ is the inactivation rate. Here we have assumed that both populations have the same diffusion constant *D*. As a consequence, mass conservation implies *n*_*i*_*(r, t)* = *n*_0_ *-n*_*a*_*(r, t*), were *n*_0_ is the total concentration of nucleators. The equation for active nucleators is supplemented with the following boundary conditions, ∂ _*r*_*n*_*a*_(*R, t*) = 0 and *D∂* _*r*_*n*_*a*_(*r*_0_, *t) =* −*qn*_*i*_*(r*_0_, *t)*. These conditions correspond to reflecting boundary conditions at the cell boundary *R* and activation of nucleators at a rate *q* at the surface of DNA, that we represent by a sphere of radius *r*_0_. The steady-state solution for the profile of active nucleators at a distance *r* from the cell center reads

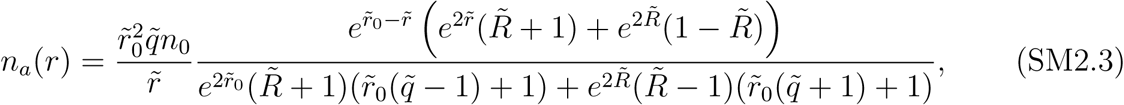

where 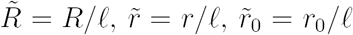, and 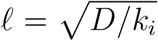 is the length scale of the diffusion and inactivation gradient. Finally, 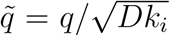 is the non-dimensional activation rate. The total number of activated nucleators in a cell of radius *R* is given by 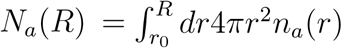, leading to

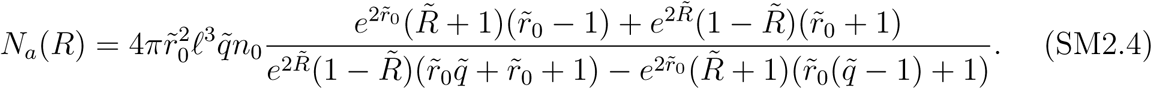

The number of active nucleators, and thus microtubules, scales like the volume of the cell for 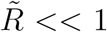, corresponding to the scaling regime, whereas it saturates for 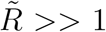, corresponding to the saturation regime. This behavior is phenomenologically captured by 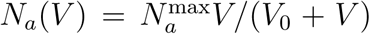, where *V* is the volume of the cell, *V*_0_ is the reaction-diffusion limited volume, and 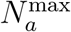 is the saturation number of active nucleators. For comparison to the full solution from equation SM2.4, see Fig S6. In the following, we will only use the phenomenological expression for the number of active nucleators. Moreover, to allow scaling behaviour other than with cell volume, we use

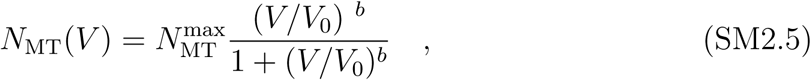

where *b* is a scaling exponent that allows for sub(super)-volume scaling. This function implies that microtubule number is constant in the large cell-regime (*V* ≫ *V*_0_), but scales with cell size in smaller cells.

### Cell boundary sequestration of an inhibitor of microtubule nucleation

Our quantitative measurements of microtubule number showed that microtubule number scales with the surface area of the cell (*b* ∼ 2*/*3) and not with cell volume. Here, we propose a model of nucleator scaling through the interplay of component limitation and membrane sequestration of an inhibitor of microtubule nucleation. We assume that a component in the cytoplasm inhibitory to microtubule nucleation acts as an enzyme to inactivate nucleators. In the well-mixed regime (*V < V*_0_), the concentration of active nucleators *n*_*a*_ depends on the balance between activation and inactivation

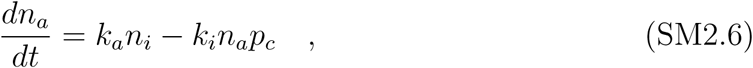

where *n*_*i*_ is the concentration of inactive nucleators, *p*_*c*_ the cytoplasmic concentration of the inhibitor, *k*_*a*_ and *k*_*i*_ are the activation and inactivation rates, respectively. The number of nucleators is conserved, such that 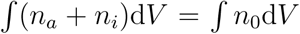, where *n*_0_ is the total nucleator concentration. We assume that the total nucleator concentration *n*_0_ is independent of cell size, which neglects changes in protein concentration that may be developmentally regulated. In the simplest case where the inhibitor does not partition into the membrane, the concentration of active nucleators *n*_*a*_ is given by

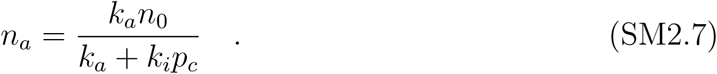

Because cytoplasmic concentrations of total nucleator *n*_0_ and inhibitor *p*_*c*_ are constant in the well-mixed regime, *n*_*a*_ is independent of cell size. Assuming that microtubule number *N*_MT_ is proportional to the number of active nucleators in the cell implies that microtubule number scales with cell volume,

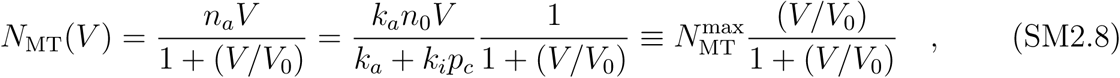

where 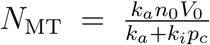. In the case where an inhibitor of microtubule nucleation is partitioned to the cell boundary, the cytoplasmic fraction of the inhibitor *p*_*c*_ in the well-mixed regime follows

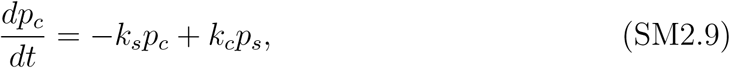

where *k*_*s*_ and *k*_*c*_ are the rate of binding and unbinding from the cell boundary, and *p*_*s*_ is the population of inhibitor in the cell boundary. Mass conservation of the total amount of inhibitor imposes 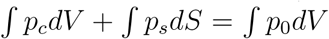, where *p*_0_ is the total inhibitor concentration. At steady state, the concentration of inhibitor *p*_*c*_ in the cytoplasm is 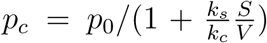. This implies that the concentration of active nucleators in the cytoplasm in the well-mixed regime is given by

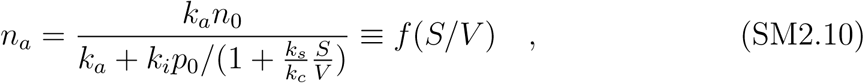

where *f*(*S/V*) is a function of the surface area-to-volume ratio. The amount of microtubules as a function of the cell volume in the full regime can thus be expressed as

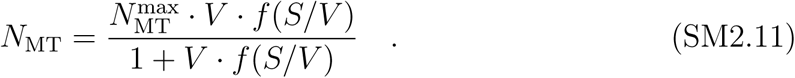

Fitting this expression of the number of microtubules to our data gives a critical radius *R*_0_ ∼ 50 *µ*m that is consistent with previous measurements (12).

### Cell size-dependent regulation of microtubule dynamics

Next, we explored the transition from constant to scaling microtubule dynamics below a critical cell size. Microtubule polymerization velocity depends on the amount of free tubulin and microtubule-associated proteins that promote microtubule growth (here referred to as MAPs) (22, 38, 39).The upper limit in microtubule polymerization velocity *v*_*p*_ is likely related to a saturation of microtubule plus ends or the microtubule lattice with MAPs (e.g. XMAP215 or EB proteins) (22, 38, 39). The observed decrease in microtubule growth velocity *v*_*p*_ as cells and spindles become smaller suggests that the number of MAPs per microtubule decreased below a critical value, such that microtubules are not saturated of MAPs anymore. Because the transition to scaling of microtubule dynamics occurs well in the spindle scaling regime–which corresponds to the well-mixed regime–, here we only consider our analysis in this scenario (*V < V*_0_). In this regime, we estimate the number of MAPs per microtubule by

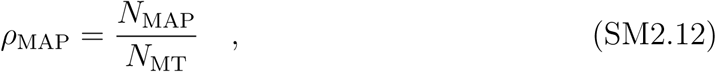

where *N*_MAP_ and *N*_MT_ are the number of MAPs and microtubules, respectively. We assume that that the number of MAPs promoting microtubule growth scale with cell volume as in the simplest limiting component model. Because microtubule number scales as sub-volume (see equation SM2.11 and Fig 4), the number of MAPs per microtubule decreases monotonically with decreasing cell size (further sequestration of components to the cell boundary that act to decrease microtubule growth leads to similar decrease in the number of MAPs per microtubule). As a consequence, when the number of MAPs per microtubule decreases below the saturation value, microtubule growth velocity will scale with cell size.

The dependence of the microtubule growth velocity *v*_*p*_ on the concentration *ρ*_MAP_ of MAPs per microtubule that takes into account their saturation can be cast into a Michaelis-Menten type equation,

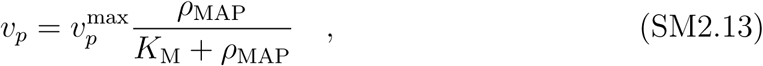

where 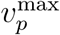 is the maximum polymerization velocity when the microtubule is saturated with MAPs and *K*_M_ is the MAP concentration at which the polymerization velocity is half of the maximum. For the concentration of MAPs per microtubule, *ρ*_MAP_, we assume that the amount of MAPs scales with cell volume (as in the simplest limiting component), and consider that the number of microtubules is given by equation SM2.11. This expression fits quantitatively the scaling of velocity polymerization with cell size (see Fig 4).

